# Evolutionary paleoecology of European rhinocerotids across the Oligocene-Miocene transition

**DOI:** 10.1101/2024.04.17.589495

**Authors:** Manon Hullot, Céline Martin, Cécile Blondel, Damien Becker, Gertrud E. Rössner

## Abstract

The Oligocene-Miocene transition witnessed great environmental and faunal changes, spanning from late Oligocene to early Miocene (MP28-MN3). Its drivers and consequences on mammals are however poorly understood. Rhinocerotoids are among the most affected taxa, reflected by great taxonomical and morphological changes. However, potential associated changes in ecology have not been explored. Here, we investigated the paleoecology of 10 rhinocerotid species coming from 15 localities across Western Europe and ranging from MP28 to MN3. We explored evolutionary trends for diet, physiology, and habitat via dental wear, hypoplasia, body mass, and stable isotopy. All rhinocerotids studied were C3 feeders, whether browsing or mixed-feeding, but clear dietary differences were observed at some localities and between Oligocene and Miocene rhinocerotids. The prevalence of hypoplasia was low (< 10 %) to moderate (< 20 %), but there were great differences by loci, species, and localities. Body mass co-variated with hypoplasia prevalence, suggesting that larger species might be more susceptible to stresses and environmental changes. We reconstructed similar warm conditions at all localities except Gaimersheim, but found greater variations in precipitation. Indeed, a clear shift in δ^13^C values was noticed at the end of the Oligocene, consistent with climatic and vegetation changes reported at that time.

## Introduction

While the Oligocene-Miocene transition (OMt) also marks the boundary between two periods (Paleogene and Neogene), this event has received little attention, hence its drivers and consequences on terrestrial faunas remain poorly understood [1]. Despite the apparent global climatic stability reconstructed during the Oligocene and Early Miocene (coolhouse episode: 34-15 Ma), major punctual (spatially and/or temporally) climatic fluctuations occurred during this interval, including the Late Oligocene Warming (26.5-24.5 Mya) and the Mi-1 glaciation event (23.28 to 22.88 Ma) [2,3]. The Late Oligocene Warming occurred between the MP26 and MP28 [3], and was responsible for a temperature increase in terrestrial environments of up to 10°C [4–6]. The Mi-1 is one of the three largest climatic aberrations of the Cenozoic, occurring at the Oligocene-Miocene boundary and responsible for a prolonged cooling lasting about 400 ky [7–9],. Both events might have critically impacted the food resources and habitats, which might be reflected in the paleoecological preferences of the fauna.

A few studies have investigated mammal occurrence during the OMt, with contrasting results. Micro-mammal occurences point towards a relative faunal homogeneity across Europe with no particular changes in communities at the Oligocene-Miocene boundary [1,10]. The trend is very different for large herbivore mammals, for which Scherler et al. [11] described a three-phased transition of the assemblages (genus and family levels) over a long period of time from the Mammal Paleogene reference level (MP) 28 to the Mammal Neogene zone (MN) 3. Moreover, some faunal changes are reported during the OMt, as classical components of Paleogene faunas disappear. This is notably the case of “hyracodontid” and amynodontid rhinocerotoids, leaving only one family of rhinoceroses from the Miocene onward: the Rhinocerotidae or true rhinoceroses.

The super-family of Rhinocerotoidea – including Eggysodontidae, Paraceratheriidae, Amynodontidae, and Rhinocerotidae – is the most abundant and diversified within the Perissodactyla [12]. More than 300 species are recognized from the Eocene onward in various terrestrial ecosystems of all continents but Antarctica and South America [13]. Rhinocerotidae arrived in Europe after the *Grande Coupure* (early Oligocene), but only reached their peak of alpha-diversity shortly after the OMt (Burdigalian) [14], with often several co-occuring species documented in the Neogene faunas. Today, rhinoceroses are the largest herbivore species with both grazing and browsing preferences [15], and the past diversity could have been even greater, as evident changes in locomotion and dental morphology are documented throughout the evolutionary history of this family [11,14]. However, rhinocerotids’ paleoecology has rarely been explored.

Here, we investigate the paleoecology of 10 rhinocerotid species coming from 15 localities across Western and Central Europe covering the Oligocene-Miocene transition interval (MP28-MN3). As climatic events might have impacted food resources and habitats, and as morphological changes have already been noted for rhinocerotids during the OMt [11], we expect to detect changes in the paleobiology and paleoecology of these taxa as well. We explore evolutionary trends for body mass (dental measurements), dietary preferences (dental wear and carbon isotopes) and stress susceptibility (enamel hypoplasia) to infer the paleoecological niche and its spatio-temporal evolution. This approach allows to discuss some aspects of niche partitioning or competition, and provides new paleoenvironmental insights (mean annual temperature and precipitation) at several localities. We anticipate a shift in dietary preferences between Oligocene and Miocene rhinocerotids and an increase of body mass during the earliest Miocene associated with the appearance of mediportal forms.

## Material and methods

We studied approximatetly 1800 teeth of rhinocerotids (see all details in ESM1) from 15 European localities dating to the OMt (MP28-MN3; Figure 1 and details on localities in ESM2). The specimens are curated at the following institutions: Bayerische Staatsammlung für Paläontologie und Geologie Munich (Gaimersheim, Pappenheim, Wintershof-West), Centre d’étude et de Conservation du Muséum Marseille (Paulhiac, Laugnac), Musée des Confluences de Lyon (Gannat), Museo Nacional de Ciencias Naturales Madrid (Loranca del Campo, Valquemado), Museum National d’Histoire Naturelle Paris (Gannat, La Milloque, Thézels), Naturhistorisches Museum Basel (Gannat, La Milloque, Laugnac, Paulhiac, Thézels, Rickenbach), Naturhistorisches Museum Bern (Engehalde, Wischberg), Naturmuseum Olten (Rickenbach), Rhinopolis (Gannat, deposited at Paleopolis), Staatliches Museum für Naturkunde Stuttgart (Tomerdingen, Ulm-Westtangente), University Claude Bernard Lyon 1 (Laugnac, Gannat), University of Poitiers (La Milloque, Thézels, Paulhiac).

**Figure 1:**
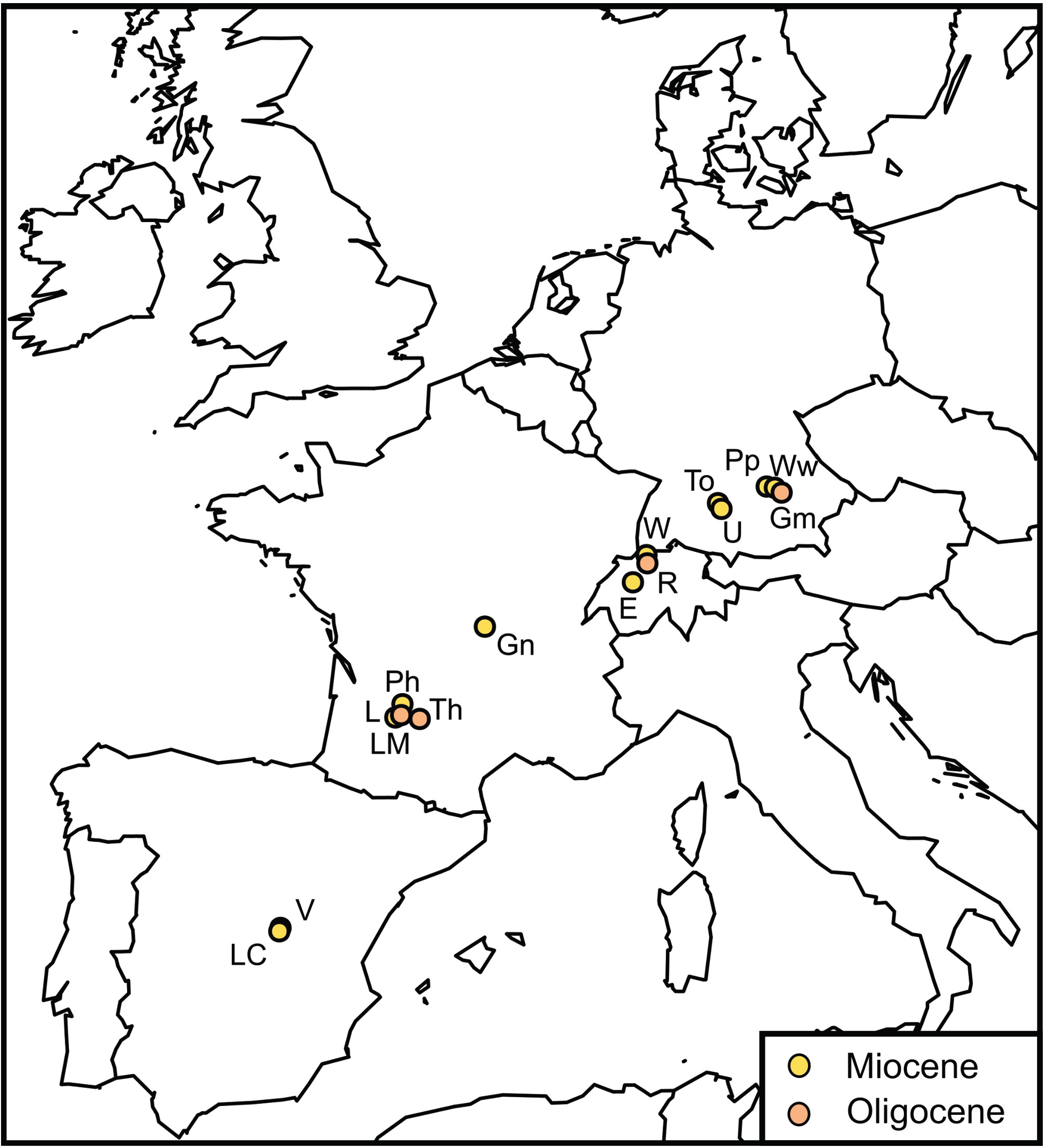
Localization of the studied localities in Central and Western Europe. Abbreviations: E – Engehalde (MN2), Gm - Gaimersheim (MP28), Gn – Gannat (MP30-MN1), L – Laugnac (MN2, reference), LC – Loranca del Campo (MN2), LM – La Milloque (MP29), Ph – Paulhiac (MN1, reference), Pp – Pappenheim (MN1), R – Rickenbach (MP29, reference), To – Tomerdingen (MN1), Th – Thézels (MP30), U – Ulm-Westtangente (MN2), V – Valquemado (MN2), W – Wischberg (MN1), Ww – Wintershof-West (MN3).

We used a multi-proxy approach to investigate paleoecology, combining stable carbon and oxygen isotopes, dental mesowear and microwear, enamel hypoplasia, and body mass estimation. The number of teeth studied for each method is detailed in Table 1 by locality and species. Each method is detailed thereafter.

**Table 1:**
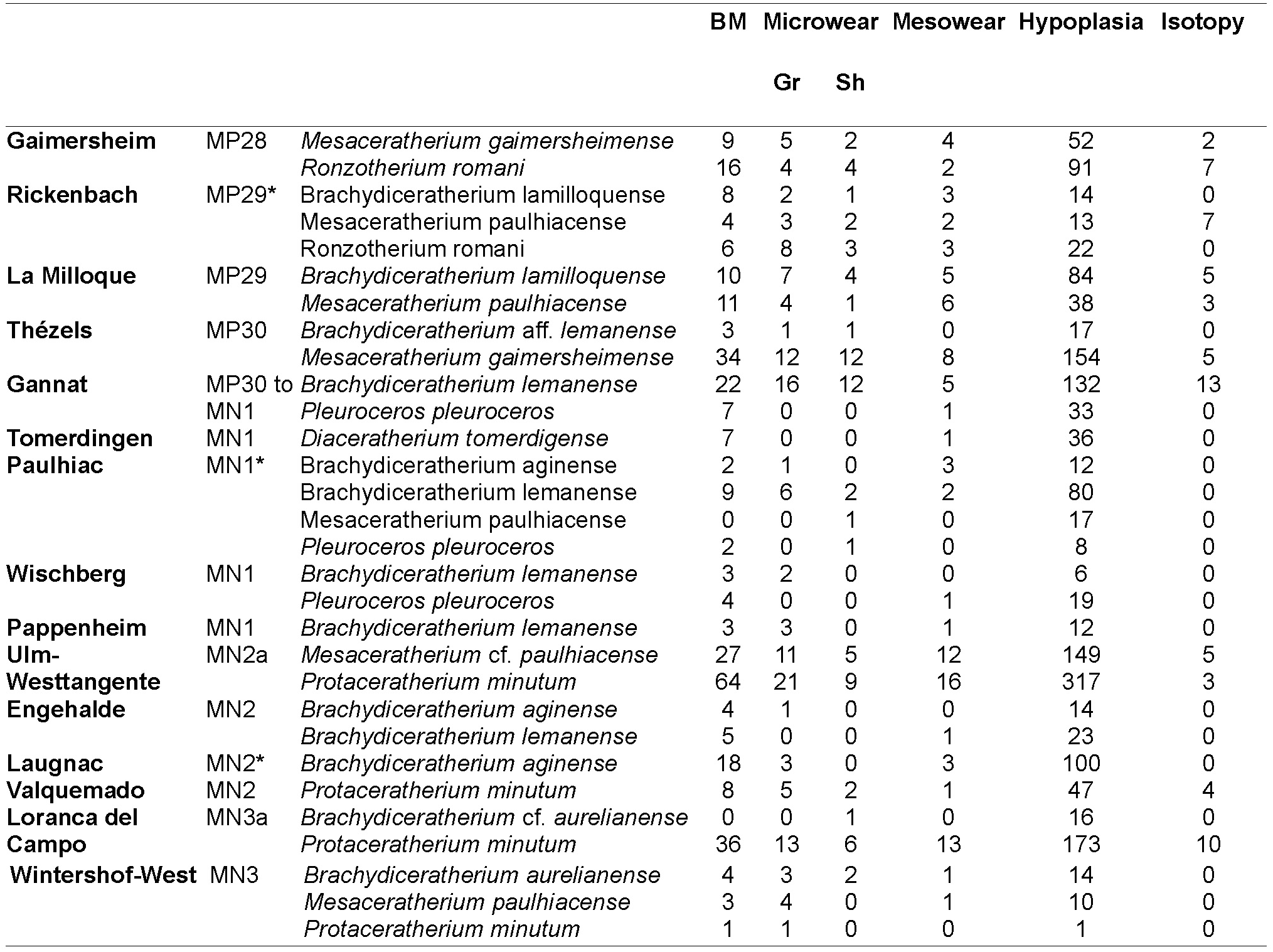
List of rhinocerotid species found at each locality studied along with the number of specimens included for each method. * indicates that the locality is the reference of the Mammal Paleogene (MP) or Neogene (MN) Zone Abbreviations: BM for body mass, Gr for grinding facet, Sh for shearing facet. Time ordering according to the following references [16–18].

### Body mass estimations

The body mass of mammals is linked to a large number of physiological and ecological traits [19,20], including metabolism rate, behavior, habitat, or spatial distribution. Although not the best proxies to estimate body mass, teeth were used here, as they are abundant and more often well preserved in the fossil record than long bones [21,22]. Classical equations linking length and width of first and second upper and lower molars were used, as detailed in Table 2.

**Table 2:**
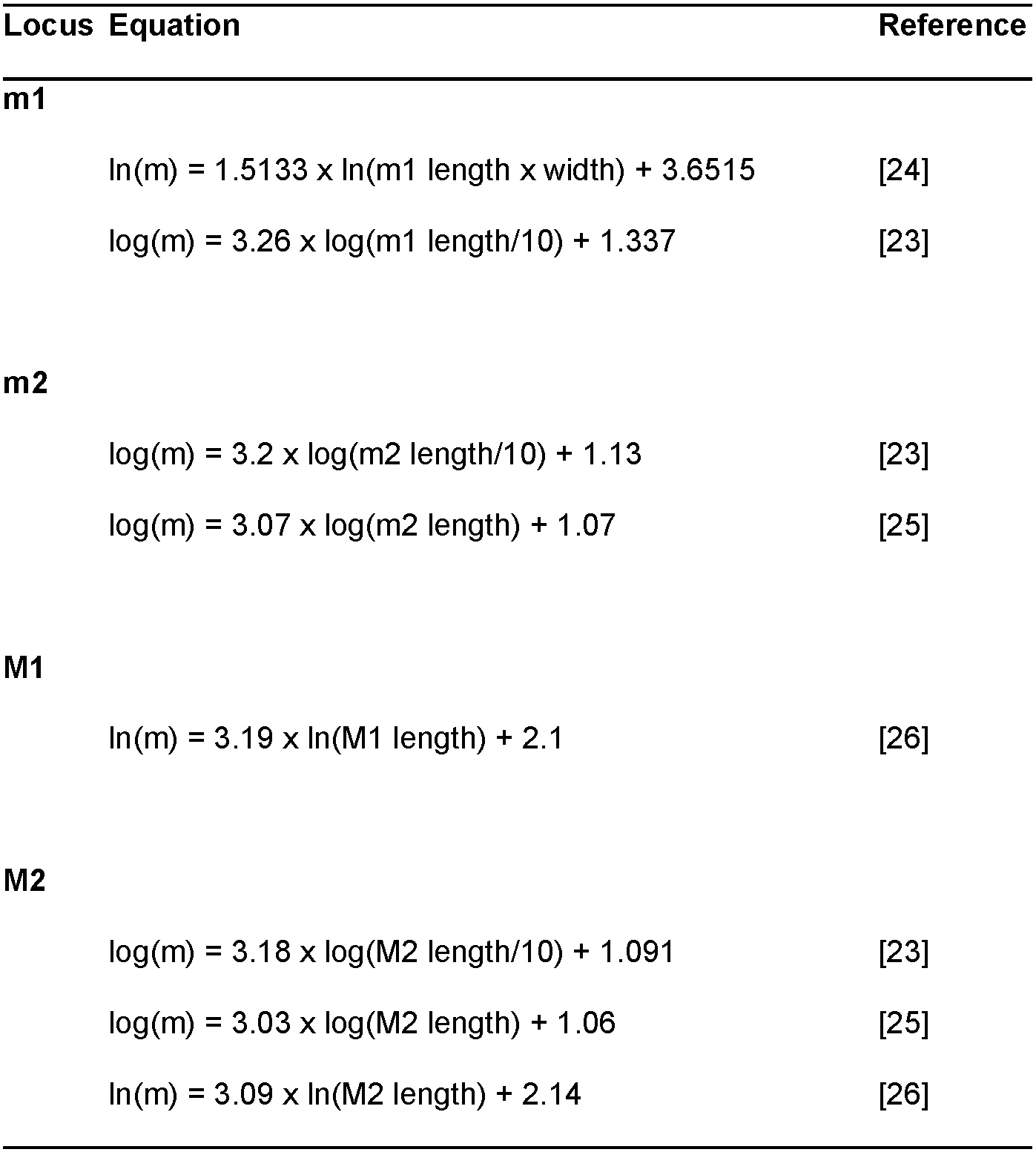
List of the dental proxies and the associated equations used in this study to estimate rhinocerotids’ body mass. Measurements are in mm for all equations and give body mass in kg for Janis [23] and in g otherwise.

### Enamel hypoplasia

Tooth enamel develop early in life and is not remodeled afterwards. Teeth may record stresses that result in developmental hiatuses and growth defects. Among these defects, enamel hypoplasia is one of the most common [27]. Enamel hypoplasias are permanent and sensitive defects, but they are individual-dependant and non-specific. They have been linked to many causes including birth [28,29], weaning [30,31], diseases [32], or nutritional stress [29,30]. No consensus nor on the method to study hypoplasia, nor on the threshold between normal and pathological enamel exist, so we chose to investigate hypoplasia with the naked-eye, as it is faster, cheaper, more used, and less prone to false positives than microscopy approaches [33]. This method consists in the macroscopical spotting and identification of the defects according to the *Fédération Dentaire Internationale* categories (linear, pit, and aplasia) and the caliper measurements of parameters (e.g., distance to enamel dentine junction, width of the defect) related to timing, severity, and duration of the defect. Other information recorded includes the number of defects, localization on the crown, and degree of severity (Figure 2).

**Figure 2:**
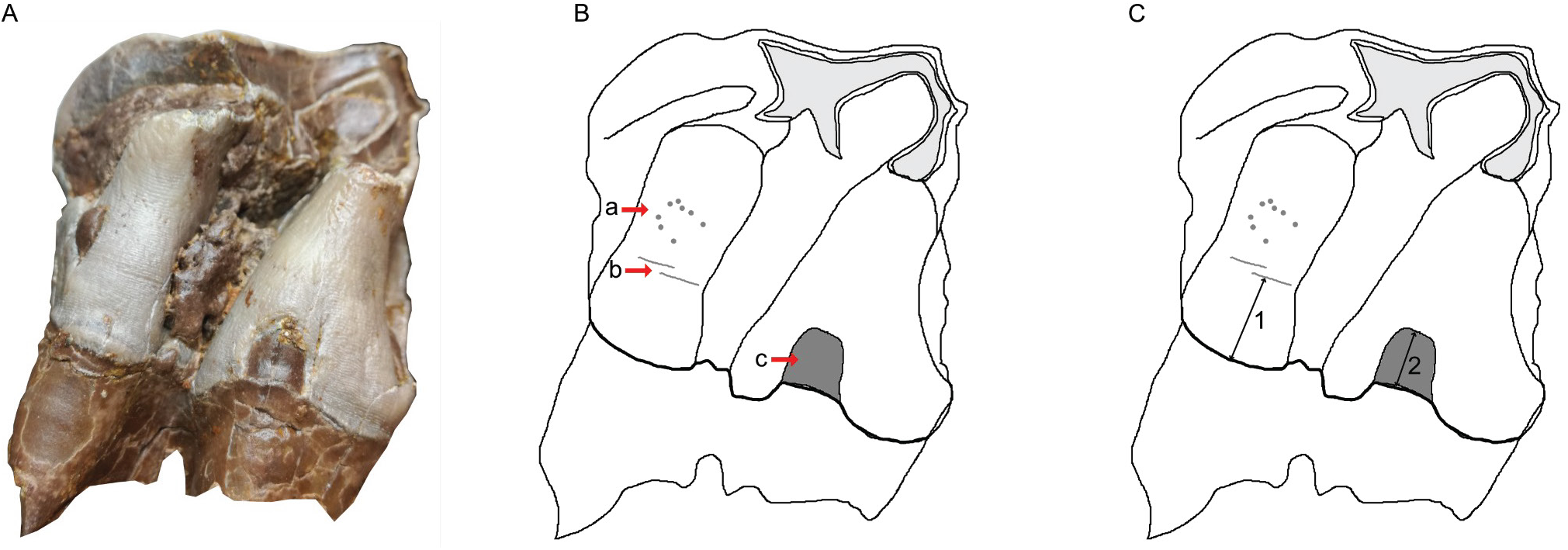
The three different types of hypoplasia considered in this study and the associated measurements. A-Lingual view of right M2 of the specimen MHNT.PAL.2004.0.58 (*H. beonense*) displaying three types of hypoplasia. B-Interpretative drawing of the photo in A illustrating the hypoplastic defects: a- pitted hypoplasia, b- linear enamel hypoplasia, and c- aplasia. C- Interpretative drawing of the photo in A illustrating the measurements: 1- distance between the base of the defect and the enamel-dentin junction, 2- width of the defect (when applicable). Figure from Hullot et al. [34].

### Carbon and oxygen stable isotopes of the carbonates in the rhinocerotids’ enamel

Carbon isotopic composition of teeth and bones is linked to the feeding behavior (C3 or C4 plants preferences) and tracks habitat openness [35,36], while oxygen isotopic composition might inform on the temperature and precipitation [36,37]. Here, we have focused on the signal from the carbonates of tooth enamel from rhinocerotids, as both isotopes can be studied at the same time (faster and cheaper).

Not all localities were sampled for isotopic analyses, due to sampling restrictions (destructive and costly), but we provided a coverage of all biostratigraphic zones of the MP28 to MN3 interval. Rhinocerotid teeth from the following localities were sampled: Gaimersheim (MP28), La Milloque (MP29), Thézels (MP30), Gannat (MP30-MN1), Ulm-Westtangente (MN2), Valquemado (MN2), and Loranca del Campo (MN3). Additionally, we included data from the literature for the locality of Rickenbach (MP29) [38]. Some specimens from La Milloque, Thézels, and Gannat were also serially sampled to investigate seasonality close to the Oligocene-Miocene boundary and the Mi-1 event.

Sampling was done on a restricted zone close to the root-crown junction (last part of the crown to develop, punctual) or along the crown (serial) on identified isolated teeth or fragments, preferably from third molars to avoid pre-weaning or weaning signal. After mechanical cleaning with a Dremmel© equipped with a diamond tip, enamel powder was sampled. As we focused on carbonates, carbon and oxygen isotopic composition could be studied at the same time, limiting the amount necessary for the analyses (between 500 and 1000 μg). Organic matter was removed following standard procedures [39] and the samples were then acidified with phosphoric acid (103 %), producing CO2 analyzed for isotopic content using a Micromass Optima Isotope Ratio Mass Spectrometer (Géosciences Montpellier). Results are expressed as ratio (‰) to the Vienna-Pee Dee Belemnite (VPDB) standard as follows:

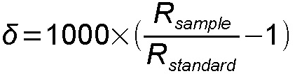 where R_sample_ refer to the ratio of 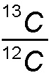 and 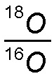 of the sample and R_standard_ to the VPDB.

The δ^13^C_diet_ can be obtained from δ^13^C_C, enamel_ taking into account the body mass and the digestive system as detailed below [40]:

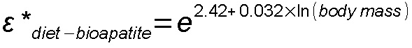

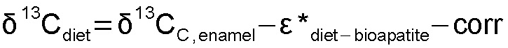 where corr is the correction factor for the variation of δ^13^C_CO2_ of the atmosphere. Post 1930, the values of δ^13^C_CO2_ are −8 ‰ [7]. Depending on the locality, the reconstructed values based on benthic foraminifera [41] are higher than today with estimates between −6.1 and −5.7 ‰. The δ^13^C_diet_ is then used to infer the mean annual precipitations (MAP) with the equation from Kohn [42]: 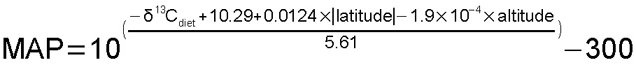 or that of Rey et al. [43]: 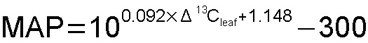 where 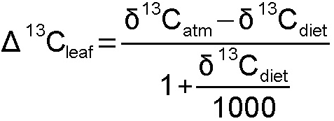.

Regarding oxygen, the δ^18^O_CO3(V-PDB)_ was converted into δ^18^O_CO3(V-SMOW)_ using the equation from Coplen et al. [44]: 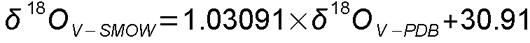. This was used to calculate the δ^18^O_precipitation_ and the mean annual temperature (MAT) detailed as follows. No reliable equation to estimate the δ^18^O_precipitation_ based on the δ^18^O_enamel_ of rhinoceros is available in the literature, so we used an equation designed for elephants [45], as their metabolism (hindgut fermenter) and size (megaherbivore) are close to that of rhinoceros [15]. The δ^18^O in the following equations are expressed in relation to the V-SMOW: 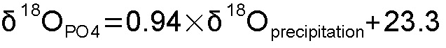 equation from Ayliffe et al. [45] for modern elephants.

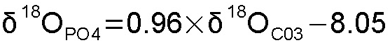 relation phosphates-carbonates from Lécuyer et al. [46].

Hence: 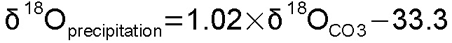

Eventually, the MAT can be calculated using the obtained δ^18^O_precipitation_:

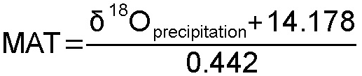 [47] or MAT=1.41×δ ^18^O_precipitation_ +23.63 [48].

### Dental wear: mesowear ruler and dental microwear texture analyses

To provide more robust dietary inferences, we combined mesowear and microwear. Both proxies give insights at different time scales: days to weeks for microwear and long-term cumulative over life for mesowear [49,50]. Mesowear is the categorization of macroscopical dental wear of labial cusp shape (relief and sharpness) into scores that can be used to infer individual dietary preferences within the classical herbivore diet categories: browser, mixed-feeder, grazer [51]. Here, we used the mesowear ruler [52], that gives scores ranging from 0 (cusp high and sharp) to 6 (cusp low and blunt; Figure 3). We only scored the paracone of upper molars (mostly M1-2, but two M3 were included) with an average wear (wear stages 4 to 7 from Hillman-Smith et al. [53], and not the sharpest cusp (metacone or paracone), as significant differences have been noted between these two cusps in rhinoceros [34,54]. With this approach, browsers have low scores (mean of extant species reported in the literature between 0-2) and grazers high ones (2.09-5.47), while mixed-feeders have intermediate values (0.4-2.74) [55]. Despite the wide use of the mean for mesowear data in the literature [50,52,56,57], we chose to use the median in this study. Indeed, the mesowear ruler yields ordinal categorical scores, which implies that the mean would assume equidistant categories or would not make sense mathematically.

**Figure 3:**
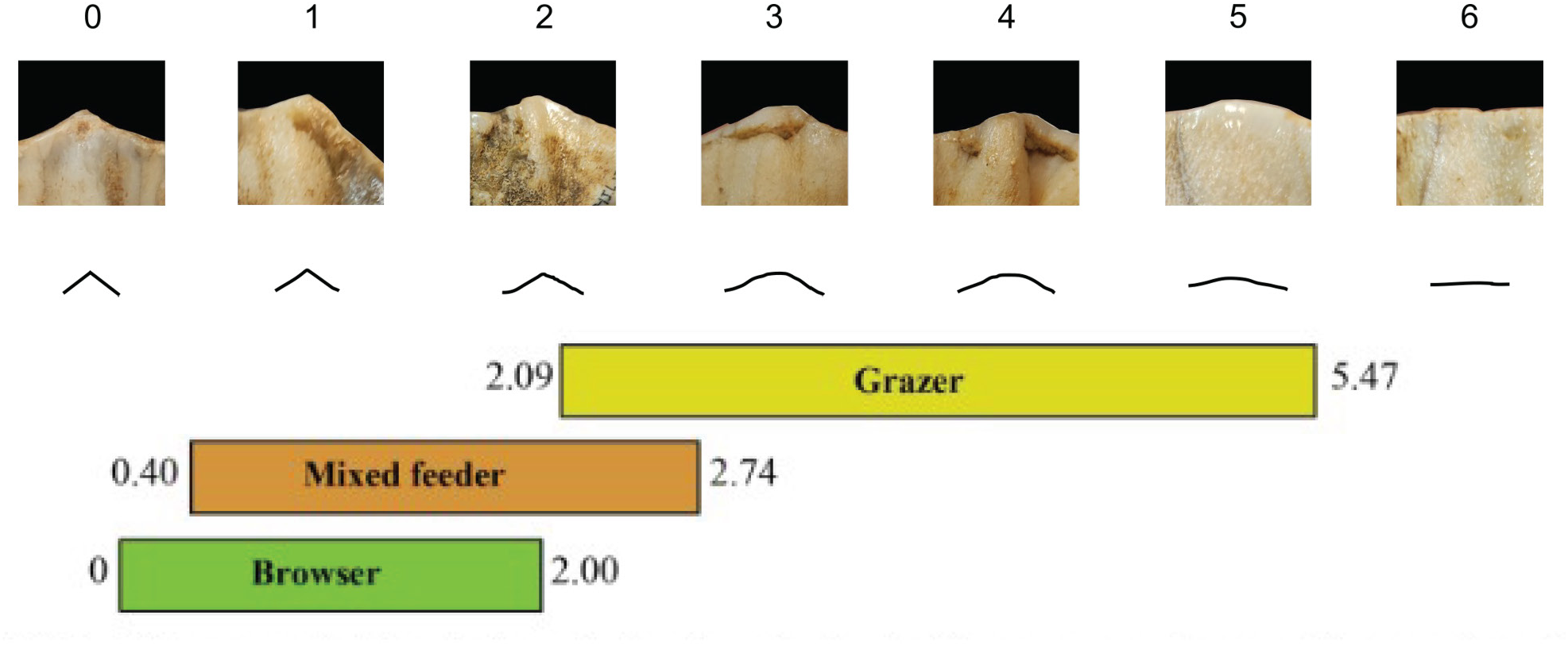
Mesowear Ruler illustrated with rhinoceros’ (*Coelodonta antiquitatis*) cusps and interpretative drawings. Modified from Jiménez-Manchón et al. [58].

Dental microwear texture analyses (DMTA) investigates tooth surface and identify wear patterns associated with the different diet categories. We followed a protocol adapted from Scott et al. [59] using scale-sensitive fractal analyses and completed the overview with ISO parameters [60]. Facets representing both phases of the mastication (grinding and shearing) were sampled on the same enamel band near the protocone, protoconid or hypoconid (Figure 4). After thorough cleaning with acetone, we molded twice well-preserved wear facets of molars (upper and lower, left and right) using dentistry silicone (Coltene Whaledent PRESIDENT The Original Regular Body ref. 60019939). The second mold was used for the analyses described hereafter.

**Figure 4:**
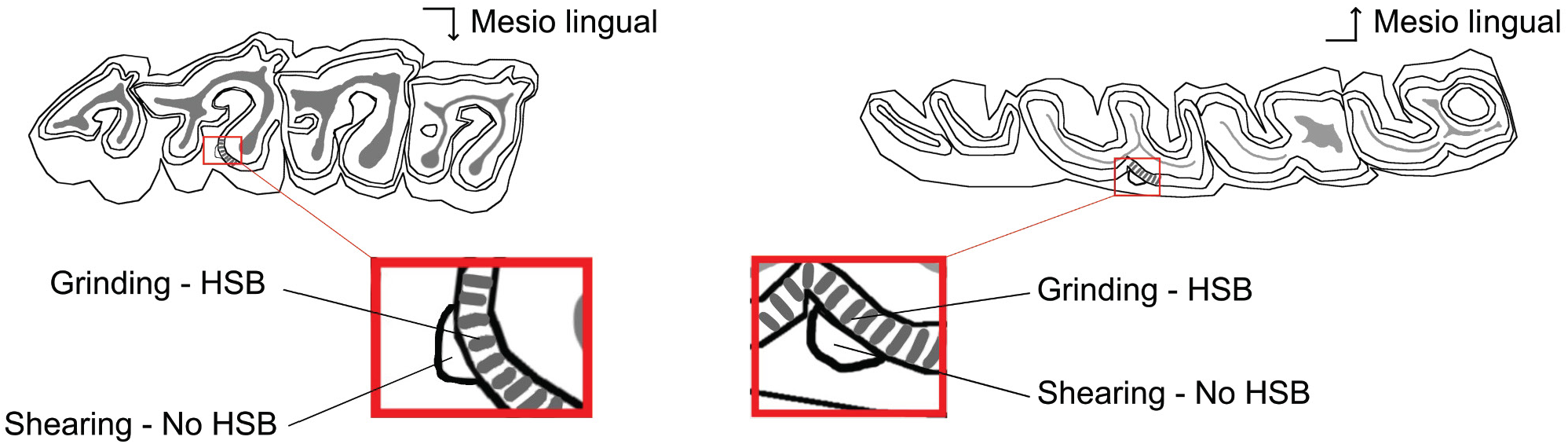
Localization of the microwear facets on rhinocerotid molars. Position of the two microwear facets (grinding and shearing) near the protocone on the second upper molar (left) and near the protoconid on second lower molar (right). Both facets are sampled on the same enamel band with (grinding) or without (shearing) Hunter-Schreger bands (HSB). Modified after Hullot et al. [61].

The molded facet was put flat under the 100× objective (Leica Microsystems; Numerical aperture: 0.90; working distance: 0.9 mm) of the Leica Map DCM8 profilometer hosted at the PALEVOPRIM Poitiers (TRIDENT), and scanned using white light confocal technology. Using LeicaMap (v.8.2; Leica Microsystems), we pre-treated the obtained scans (.plu files) as follows: inversion of the surface (as they are negative replicas of the actual surface), replacement of the non-measured points (less than 1%) by the mean of the neighboring points, removal of aberrant peaks [62], leveling of the surface, removal of form (polynomial of degree 8) and selection of a 200 × 200 μm area (1551 × 1551 pixels) saved as a digital elevation model (.sur) to be further analyzed. Textural and ISO parameters of the selected surfaces (200 × 200 μm) were then estimated in MountainsMaps® (v.8.2). Our study will focus on the following texture variables, described in details in Scott et al. [63]:

- anisotropy measures the orientation concentration of surface roughness. Several parameters can indicate the anisotropy of the surface:

- epLsar (exact proportion of length-scale anisotropy of relief) or NewepLsar. The latter is the corrected value of epLsar in MountainsMaps® compared to Toothfrax (software previously used for DMTA but not supported anymore) as there was an error in the code to calculate this parameter in Toothfrax [64];
- Str is a spatial parameter of the international standard ISO 25178 (specification and measurement of 3D surface textures). It is the ratio of Rmin/Rmax, where Rmin and Rmax are respectively the minor and major axes of the intersection ellipse between the plane z = s with the autocorrelation function f_ACF_(tx, ty). Rmin is the autocorrelation length, i.e. the horizontal distance of the f_ACF_(tx, ty) which decays fastest to a specified value s between 0 and 1. Here, we considered s = 0.5. Low values of Str indicate strong anisotropy;
- complexity or area-scale fractal complexity (Asfc) evaluates the roughness at a given scale;
- heterogeneity of the complexity (Hasfc) investigates the variation of complexity at a given scale (here 3 × 3 and 9 × 9) within the studied 200 × 200 μm area.

The interpretation of DMTA signatures in fossil specimens, is based here on the values and thresholds proposed in extant species [61,65]. Moreover, we used a dataset of extant species of rhinoceros and tapirs with their associated inferred diets [61,66]. The dataset consists of 17 specimens of *Ceratotherium simum*, four of *Dicerorhinus sumatrensis*, 21 of *Diceros bicornis*, 15 of *Rhinoceros sondaicus* (one new specimen), five of *Rhinoceros unicornis* (one new specimen), and 15 of *Tapirus terrestris*. Results of DMTA for these species are available in ESM2 (Fig 1).

### Statistics and figures

Statistics and figures were done in R (v 4.2.3) equipped with ggplot2 [67], cowplot [68], gridExtra [69]. Inkscape (v 1.0.1) was also used for figures. We favored the use of non-parametric tests, due to limited sample size at some localities, which prevents from testing if the distribution of the data is normal (Table 1). For microwear data however, we used a Box-Cox transformation to apply the parametric MANOVA and ANOVA. Following the recent statement of the American Statistical Association (ASA) on p-values [70], we favored giving exact values and we tried to be critical regarding the classical thresholds of “statistical significativity”.

## Results

### Body mass estimations

The sample studied includes six genera of rhinocerotids. All the species of three genera – *Ronzotherium*, *Brachydiceratherium*, and *Diaceratherium –* were large-sized rhinoceros, exceeding the megaherbivore threshold of 1000 kg [15]. For *Ronzotherium romani*, the mean body mass was estimated between 1800-2400 kg (Table 3). Regarding the two teleoceratines, the mean body mass estimates of *Brachydiceratherium* spp. range from 1000 to 2000 kg depending on the locality and the species, whereas *Diaceratherium tomerdigense* (monotypic genus, found only in Tomerdingen) reached about 1500 kg (Table 3).

**Table 3:**
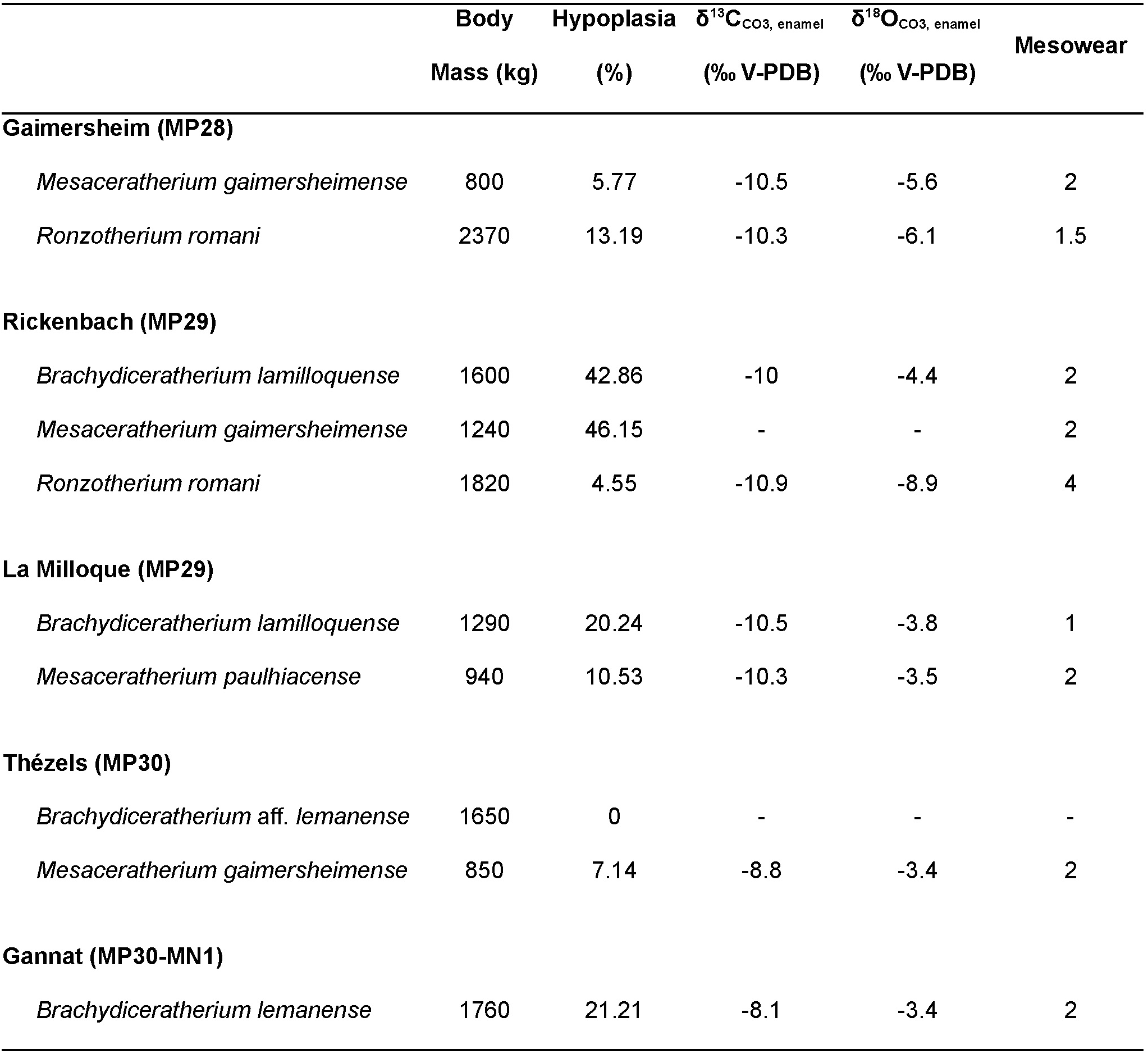
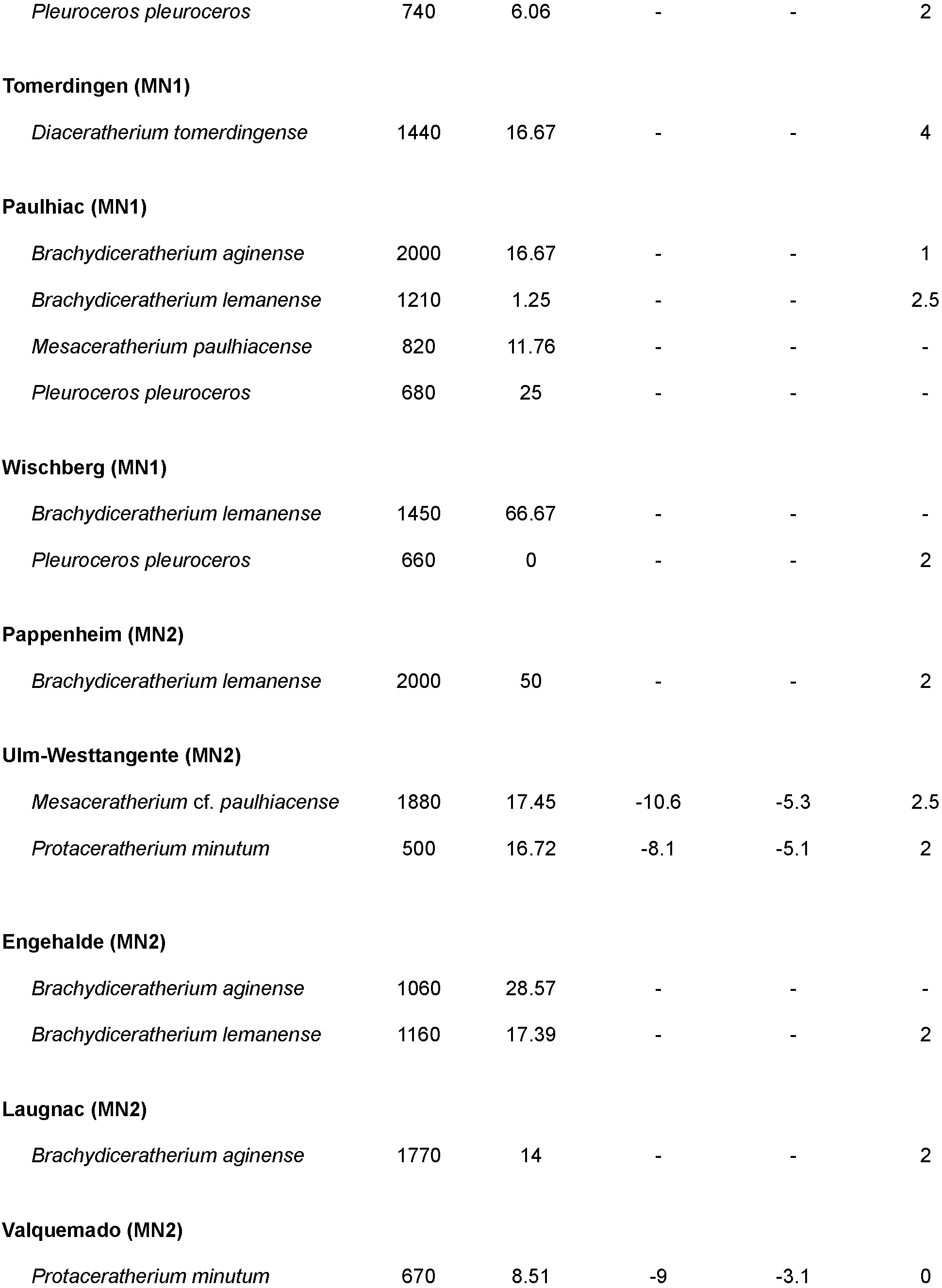
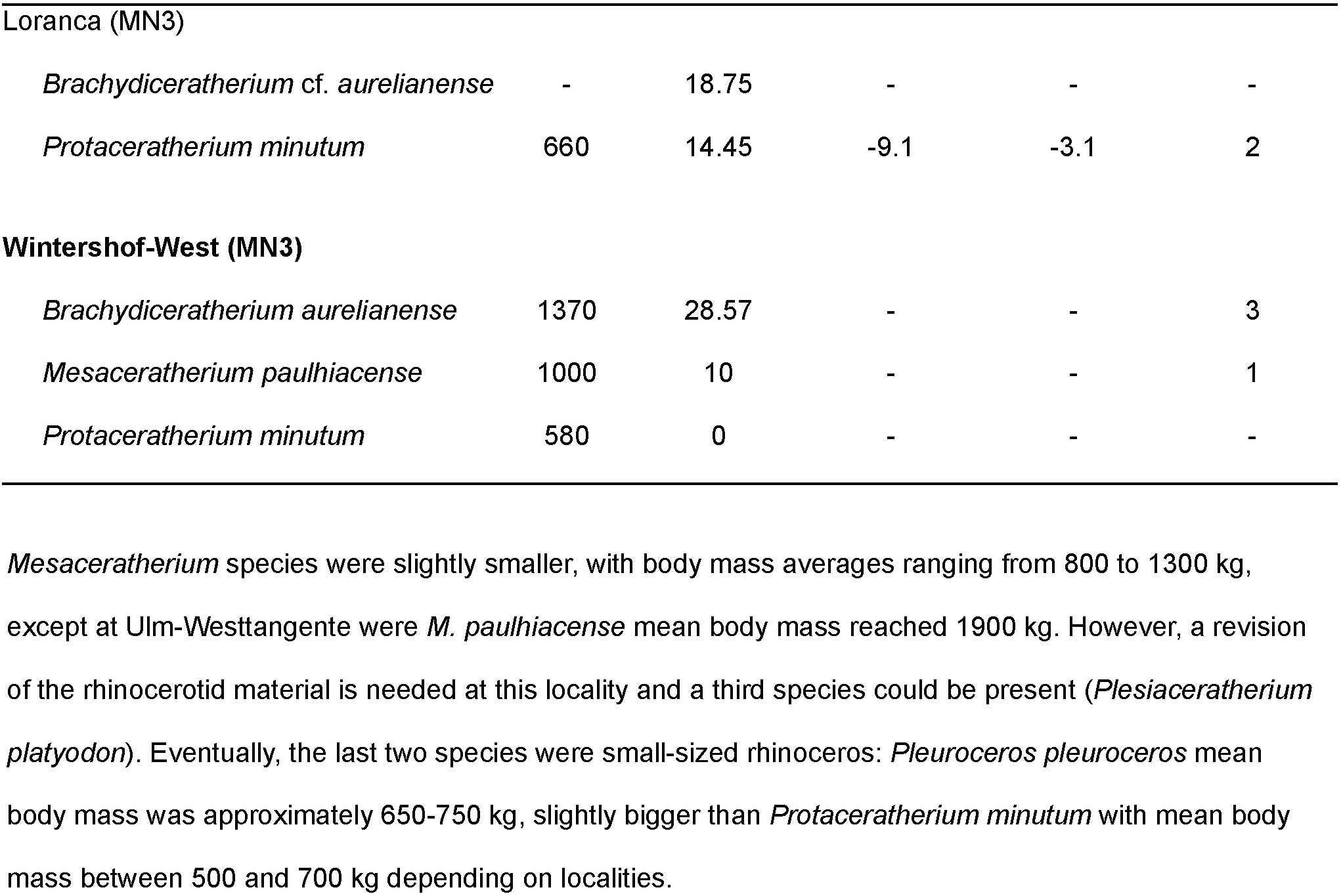
Summary of the results for body mass, enamel hypoplasia, stable isotopy (carbon and oxygen) and mesowear for the rhinocerotids of the Oligocene-Miocene transition. Mean (median for mesowear as it is categorical) by locality and by species for each parameter.

| | Body<br>Mass (kg) | Hypoplasia<br>(%) | $\delta^{13}\text{C}_{\text{CO}_3, \text{enamel}}$<br>(‰ V-PDB) | $\delta^{18}\text{O}_{\text{CO}_3, \text{enamel}}$<br>(‰ V-PDB) | Mesowear |
| --- | --- | --- | --- | --- | --- |
| **Gaimersheim (MP28)** |  |  |  |  |  |
| *Mesaceratherium gaimersheimense* | 800 | 5.77 | -10.5 | -5.6 | 2 |
| *Ronzotherium romani* | 2370 | 13.19 | -10.3 | -6.1 | 1.5 |
| **Rickenbach (MP29)** |  |  |  |  |  |
| *Brachydiceratherium lamilloquense* | 1600 | 42.86 | -10 | -4.4 | 2 |
| *Mesaceratherium gaimersheimense* | 1240 | 46.15 | - | - | 2 |
| *Ronzotherium romani* | 1820 | 4.55 | -10.9 | -8.9 | 4 |
| **La Milloque (MP29)** |  |  |  |  |  |
| *Brachydiceratherium lamilloquense* | 1290 | 20.24 | -10.5 | -3.8 | 1 |
| *Mesaceratherium paulhiacense* | 940 | 10.53 | -10.3 | -3.5 | 2 |
| **Thézels (MP30)** |  |  |  |  |  |
| *Brachydiceratherium* aff. *lemanense* | 1650 | 0 | - | - | - |
| *Mesaceratherium gaimersheimense* | 850 | 7.14 | -8.8 | -3.4 | 2 |
| **Gannat (MP30-MN1)** |  |  |  |  |  |
| *Brachydiceratherium lemanense* | 1760 | 21.21 | -8.1 | -3.4 | 2 |
| *Pleuroceros pleuroceros* | 740 | 6.06 | - | - | 2 |
| **Tomerdingen (MN1)** |  |  |  |  |  |
| *Diaceratherium tomerdingense* | 1440 | 16.67 | - | - | 4 |
| **Paulhiac (MN1)** |  |  |  |  |  |
| *Brachydiceratherium aginense* | 2000 | 16.67 | - | - | 1 |
| *Brachydiceratherium lemanense* | 1210 | 1.25 | - | - | 2.5 |
| *Mesaceratherium paulhiacense* | 820 | 11.76 | - | - | - |
| *Pleuroceros pleuroceros* | 680 | 25 | - | - | - |
| **Wischberg (MN1)** |  |  |  |  |  |
| *Brachydiceratherium lemanense* | 1450 | 66.67 | - | - | - |
| *Pleuroceros pleuroceros* | 660 | 0 | - | - | 2 |
| **Pappenheim (MN2)** |  |  |  |  |  |
| *Brachydiceratherium lemanense* | 2000 | 50 | - | - | 2 |
| **Ulm-Westtangente (MN2)** |  |  |  |  |  |
| *Mesaceratherium cf. paulhiacense* | 1880 | 17.45 | -10.6 | -5.3 | 2.5 |
| *Protaceratherium minutum* | 500 | 16.72 | -8.1 | -5.1 | 2 |
| **Engelhalde (MN2)** |  |  |  |  |  |
| *Brachydiceratherium aginense* | 1060 | 28.57 | - | - | - |
| *Brachydiceratherium lemanense* | 1160 | 17.39 | - | - | 2 |
| **Laugnac (MN2)** |  |  |  |  |  |
| *Brachydiceratherium aginense* | 1770 | 14 | - | - | 2 |
| **Valquemado (MN2)** |  |  |  |  |  |
| *Protaceratherium minutum* | 670 | 8.51 | -9 | -3.1 | 0 |
| Loranca (MN3) |  |  |  |  |  |
| *Brachydiceratherium* cf. *aurelianense* | - | 18.75 | - | - | - |
| *Protaceratherium minutum* | 660 | 14.45 | -9.1 | -3.1 | 2 |
| Wintershof-West (MN3) |  |  |  |  |  |
| *Brachydiceratherium aurelianense* | 1370 | 28.57 | - | - | 3 |
| *Mesaceratherium paulhiacense* | 1000 | 10 | - | - | 1 |
| *Protaceratherium minutum* | 580 | 0 | - | - | - |

### Hypoplasia prevalence

In total, 251 teeth out of the 1704 (14.73 %) examined for hypoplasia showed at least one defect, representing a rather moderate global prevalence of hypoplasia during the Late Oligocene – Early Miocene interval. There were however clear differences between species, localities and time intervals (3; Figure 5). Rhinocerotids from Oligocene localities had a lower global prevalence of hypoplasia (12.37 %; 60/485 teeth) than the Miocene ones (15.28 %; 161/1054), but the difference was not statistically significant (Pearson’s Chi-squared test X-squared = 4.0618, df = 2, p-value = 0.1312; ESM1). When considering the biostratigraphical units (MP and MN zones), some pairs had low p-values (between 0.0594 and 0.005) when compared with a pairwise Wilcoxon test (ESM1). This revealed time intervals of low hypoplasia prevalence (< 11 %) during MP28, MP30, and MN1, and intervals of higher prevalence (between 15 % and 20 %) during MP29, MN2, and MN3. The different zones contained however various numbers of specimens and localities (only one locality: MP28 – Gaimersheim, MP30 – Thézels; MN2 dominated by Ulm-Westtangente).

**Figure 5:**
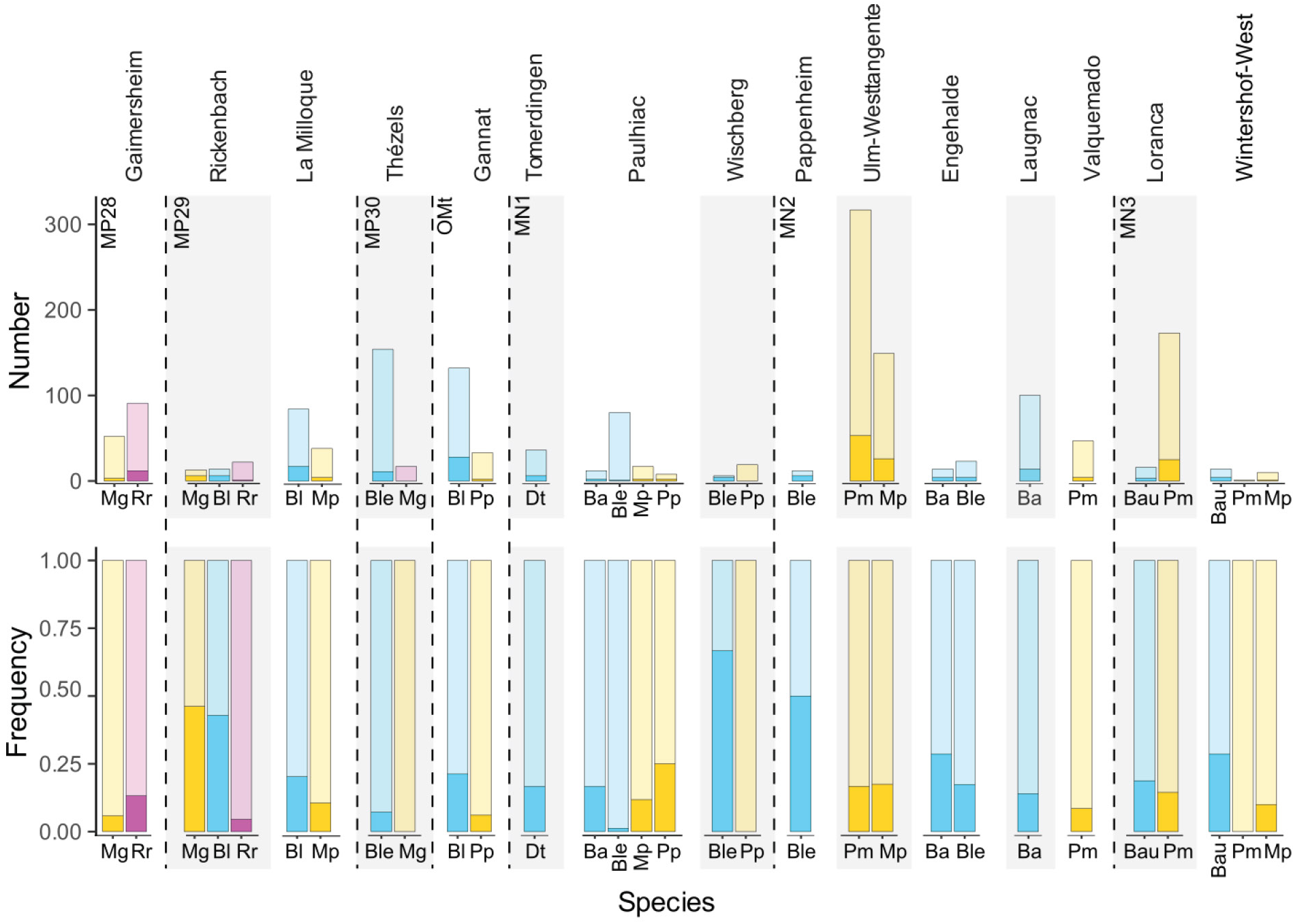
Hypoplasia prevalence by rhinocerotid species and locality of the Oligocene-Miocene transition. Abbreviations: Ba – *Brachydiceratherium aginense*, Bau – *B. aurelianense*, Bl – *B. lamilloquense*, Ble – *B. lemanense*, Dt – *Diaceratherium tomerdingense*, Mg – *Mesaceratherium gaimersheimense*, Mp – *M. paulhiacense*, Pm – *Protaceratherium minutum*, Pp – *Pleuroceros pleuroceros*, Rr – *Ronzotherium romani*. Color code: blue – Teleoceratiina, yellow – aceratheres *sensu* lato, pink – basal stem rhinocerotids Dark shades indicate hypoplastic teeth, while light shades show unaffected ones.

Concerning genus, *Brachydiceratherium* was the most affected with about 17.75 % (93/524) of studied teeth having hypoplasias. *Brachydiceratherium* was also the most common genus, present at nearly all localities. In contrast, *Pleuroceros* had the lowest frequency of hypoplasias (4/60 = 6.67 %), but this genus was only found at two localities (Gannat and Wischberg) and totaling 60 teeth. Eventually, the other genera (*Diaceratherium*, *Ronzotherium*, *Mesaceratherium, Protaceratherium*) had similar hypoplasia prevalence (pairwise Wilcoxon, p-values > 0.22; ESM1), oscillating between 12 and 17 % of hypoplastic teeth.

Regarding locality, the prevalence of hypoplasia was moderate (> 10 %) to high (> 20 %) at all of them, except at Valquemado (4/47, 8.51 %), Thézels (11/171, 6.43 %), and Paulhiac (7/117, 5.98 %) where it was low. The rhinocerotids from Pappenheim (6/12 teeth, 50 %), Rickenbach (13/49, 26.53 %), and Engehalde (8/37, 21.62 %) were the most affected, but the sample sizes were much smaller. A pairwise Wilcoxon test confirmed differences between most pairs of least affected vs. most affected localities (p-values < 0,05; ESM1).

### Enamel stable isotopy (carbon and oxygen)

All specimens were in the range of C3 feeding, with values of the δ^13^C_diet_ comprised between −27.9 and −23.1 ‰. There was a clear shift towards higher values of δ^13^C_diet_ for rhinocerotoids between La Milloque (MP29) and Thézels (MP30; Figure 6). The cutoff for this shift is around −25.6 ‰, very closed to the modern biotopes threshold between woodland-mesic C3 grassland and open woodland-xeric C3 grassland (−25 ‰; Figure 6). All specimens from Gaimersheim, Rickenbach, and La Milloque had values of δ^13^C_diet_ below this biotopes threshold, whereas most specimens of Loranca and Thézels had values close to −25 ‰ or slightly above (± 0.5 ‰). The specimens from Valquemado (*P. minutum*, n = 4), Gannat (*B. lemanense*, n= 13) and Ulm-Westtangente were on both sides of the threshold (low values for *M. paulhiacense*, n = 4; high values for *P. minutum*, n = 3). The lowest values of δ^13^C_diet_ are mostly specimens of *B. lamilloquense* and *M. paulhiacense*, whereas the highest values are specimens of *P. minutum* and *B. lemanense*.

**Figure 6:**
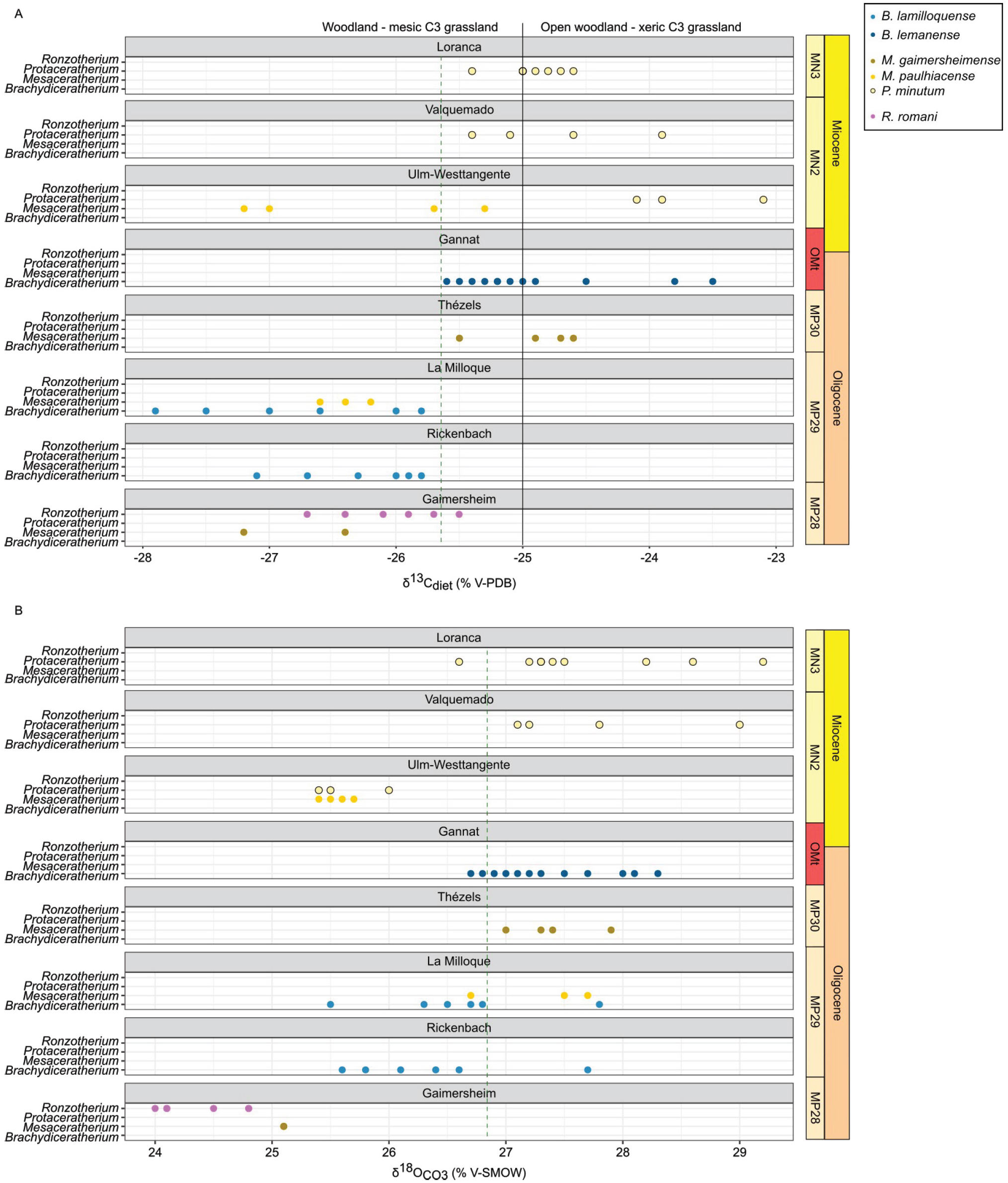
Carbon (A) and oxygen (B) content of the diet and enamel of rhinocerotids from various late Oligocene to early Miocene localities. Green dashed lines: shift, black lines: cutoffs between different biomes. Color code by species as detailed in the figure.

Regarding the oxygen content, values of the δ^18^O_CO3, SMOW_ ranged from 24 to 29.2 ‰. All specimens from Gaimersheim (*R. romani*, n = 6; *M. gaimersheimense*, n = 2), clustered together and displayed the lowest range of values (δ^18^O_CO3, SMOW_ ≤ 25.1 ‰), whereas the ones from La Milloque (*B. lamilloquense*, n = 6; *M. paulhiacense* n = 3) took a wide range of values from 25.5 to 27.8 ‰. Similarly to the δ^13^C_diet_, there is a chronological shift in the values of δ^18^O_CO3, SMOW_ between La Milloque and Thézels. The values at Ulm-Westtangente (MN2) are however more similar to the Oligocene ones.

### Dental wear: mesowear and microwear

Depending on the locality and the species, the median mesowear score was between 0 (*P. minutum* at Valquemado, n = 1) and 4 (*D. tomerdingense* at Tomerdingen, n = 1 and *R. romani* at Rickenbach, n = 3). About half of the median values by species and localities (13/24, Table 3) were however equal to 2, suggesting browsing or mixed feeding habits. The sample size of each species at each locality was often small (n < 5) or null due to preservation issues (Table 1), which limited the statistical power to compare localities and species’ spatio-temporal evolution.

Wilcoxon test revealed differences in mesowear scores at the species and genus levels. Pairwise tests failed to detect clear differences between species (p-values > 0.18), but highlighted some at the genus level (probably due to greater sample size), between *Mesaceratherium* and *Brachydiceratherium* (p-value = 0.021), and between *Mesaceratherium* and *Protaceratherium* (p-value = 0.057). Indeed, *Mesaceratherium* had greater mesowear scores than both other genera, suggesting more abrasive dietary preferences for *Mesaceratherium* specimens. There was however little changes by species between the different time intervals and localities (Table 3, ESM1), indicating limited evidence for dietary changes over space and time within our sample.

The DMTA revealed more differences, although most of the sampled fossil rhinocerotids fell in the range of extant browsers or mixed-feeders. We conducted a MANOVA on DMTA variables (Asfc, NewepLsar, H9, H81, and Str) by facet, species, and locality (parameters). All parameters had a marked influence (p-values < 10^-4^) on the DMT signature observed. To precise the differences, we ran an ANOVA for each variable. Both H9 a,d H81 had no difference for any parameters nor interaction of parameters (e.g., species * locality).

Regarding Asfc, all parameters showed differences (p-values < 0.02). In the case of NewepLsar only facet had a marked influence (p-value = 0.00132). Eventually, for Str, species (p-value = 3.67 x 10^-16^), locality (p-value = 1.09 x 10^-8^) and the interaction of species with locality (p-value = 0.051) all had a marked influence. As locality and species had more than two states, we used LSD post-hoc tests to investigate the differences highlighted by the ANOVAs for Asfc and Str.

The LSD post hocs (ESM1) did not reveal specific pairs with differences regarding Asfc for locality and only one for species (*P. minutum* and *B. lemanense*; p-value = 0,0111). Concerning Str, several pairs had low p-values for both locality and species. Most species differences were attributed to *M. gaimersheimense*, that had lower Str values than all other species studied (p-values < 0.0393), except *B. aurelianense*. Some differences were also observed for *B. lamilloquense* (high Str values) compared to *P. minutum* (p-value = 7.89 x 10^-6^), *B. lemanense* (p-value = 0.0278), and *M. paulhiacense* (0.0713), and for *P. minutum* (low Str values) compared to *B. aginense* (p-value = 0.0084; ESM1). Similarly, all differences for locality were found in pairs containing either Rickenbach (higher Str values) and/or Thézels (lower Str values).

As Asfc and Str were the two parameters with the most variations for species and locality, we plotted them (Figure 7). Contrary to mesowear, DMT signature changed over time and space for some species (see also ESM2, Fig 2 for a plot by species). Major changes of dietary preferences were observed for *B. lemanense*, which shows a decrease of anisotropy (inverse of Str) and increase of complexity (Asfc) between the MP30 (Thézels, grinding: mean Str = 0.12, mean Asfc = 1.23) and the MN2 (Pappenheim, grinding: mean Str = 0.73, mean Asfc = 2.01), except at Wischberg (grinding: mean Str = 0.34, mean Asfc = 1.14). Important variations of DMT are also found for *M. gaimersheimense,* that oscillates from a mixed-feeder profile at Gaimersheim, browser at Rickenbach (grinding: mean Str = 0.80, mean Asfc = 2.31) and to a very abrasive diet at Thézels (grinding: mean Str = 0.14, mean Asfc = 1.18). More subtle variations were found for *M. paulhiacense* with a more complex and anisotrope texture at Wintershof-West (grinding: mean Str = 0.41, mean Asfc = 2.29) than at La Milloque (grinding: mean Str = 0.57, mean Asfc = 1.90) and Ulm-Westtangente (grinding: mean Str = 0.56, mean Asfc = 1.32), for which the signatures are relatively similar. Eventually, *P. minutum* have slightly higher complexity and lower anisotropy at Iberian localities (Valquemado, grinding: mean Str = 0.51, mean Asfc = 2.21; Loranca del Campo, grinding: mean Str = 0.45, mean Asfc = 1.98) than at Ulm-Westtangente.

**Figure 7:**
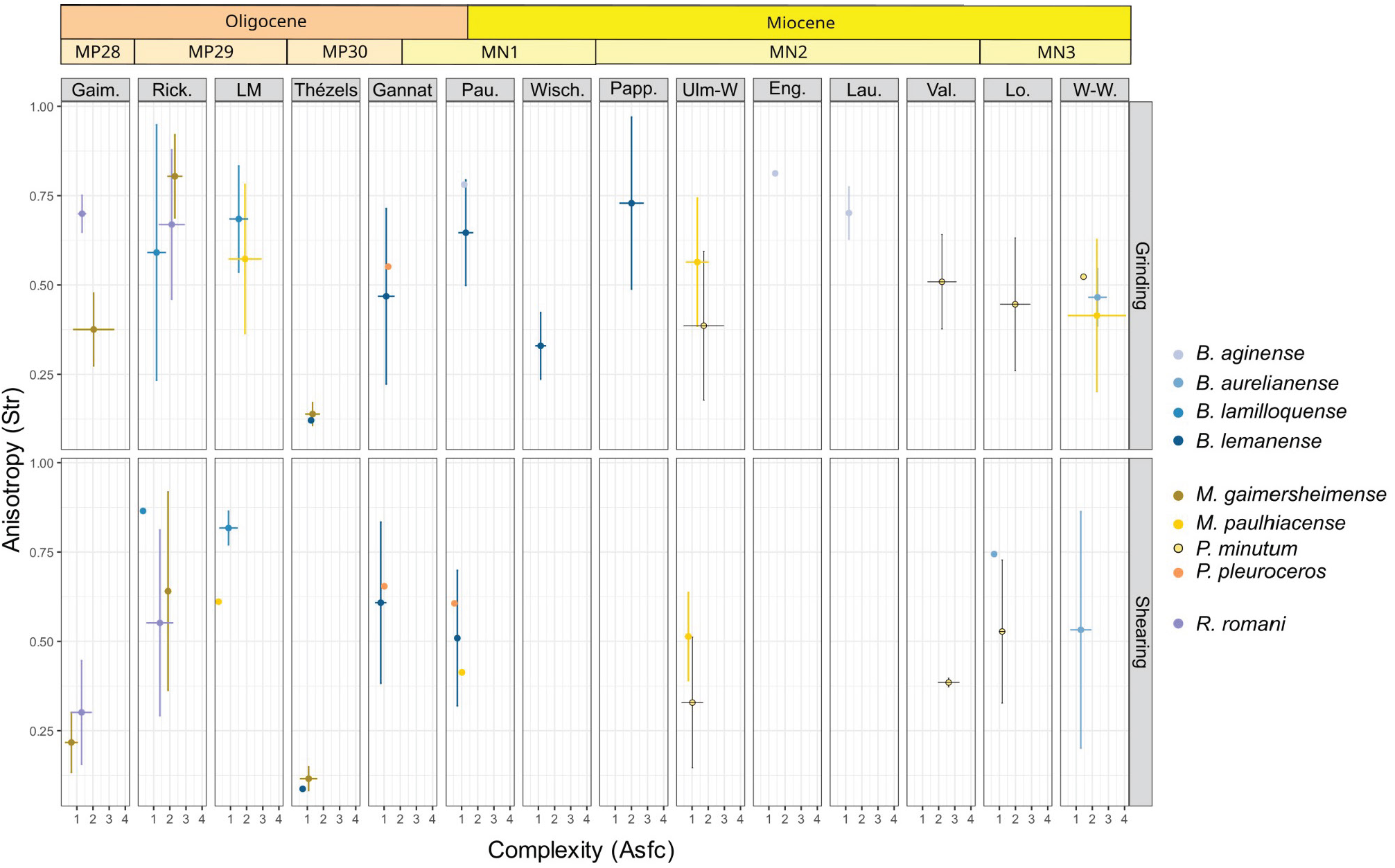
Anisotropy (Str) against complexity (Asfc) by locality and species. Localities ordered chronologically from the oldest (left) to the youngest (right): Gaimersheim (MP28), Rickenbach (MP29), La Milloque (MP29), Thézels (MP30), Gannat (MP30-MN1), Paulhiac (MN1), Wischberg (MN1), Pappenheim (MN2), Ulm-Westtangente (MN2), Engehalde (MN2), Laugnac (MN2), Valquemado (MN2), Loranca del Campo (MN3), Wintershof-West (MN3). Color code by species as detailed in the figure.

### General results and evolutionary trends

Body mass, mesowear, and hypoplasia prevalence were tracked by species and locality and plotted alongside (Figure 8, mesowear in Fig 3 ESM2). Similar trends were observed for all parameters, notably body mass and hypoplasia (rho = 0.3365995, S = 2982, p-value = 0.06895), but depended on the species. Teleoceratines and aceratheres *sensu lato* were the most common rhinocerotids in the dataset, found at nearly all localities and allowing temporal tracking of the variations. *Brachydiceratherium lamilloquense* is the oldest species of teleoceratines in the dataset, present only during MP29 at Rickenbach and La Milloque. Body mass (1600 → 1290 kg), prevalence of hypoplasia (42.85 → 20.24 %), and median mesowear ruler (2 → 1) all decrease between Rickenbach and the slightly younger La Milloque (Figure 8, Fig 3 ESM2). During the latest Oligocene and early Miocene (MP30 – MN2), *B. lamilloquense* is replaced by *B. lemanense*. This second species is documented at many localities (Thézels, Gannat, Paulhiac, Wischberg, Pappenheim, Engehalde). The median mesowear ruler scores remain similar at 2 during the interval except at Paulhiac (2,5), but greater variation of body mass and hypoplasia prevalence are noted (Figure 8). First, there is an increase during late Oligocene (Thézels: 1650 kg, 0 %→ Gannat: 1760 kg, 21.21 %), then a decrease during the earliest Miocene (Paulhiac: 1210 kg, 1.25 %), followed by a peak at Wischberg (1450 kg, 66.67 %) and a final decrease during MN2 (Engehalde: 1160 kg, 17.39 %). Another species was also found during the earliest Miocene (Aquitanian: MN1 – MN2): *B. aginense*. For this species, different trends are observed for the three parameters: body mass drops at Engehalde (1060 kg vs. 2000 kg at Paulhiac [older] and 1770 kg at Laugnac [younger]) while hypoplasia prevalence peaks at this same locality (28.57 % vs. 16.67 % at Paulhiac and 14 % at Laugnac), and mesowear increases between Paulhiac and Laugnac (1 vs. 2 respectively). Eventually, *B. aurelianense* is found during the MN3. Body mass (1370 kg) and mesowear ruler (3) were only available at Wintershof-West, while hypoplasia was recorded at both MN3 locality (18.75 % at Loranca and 28.57 % at Wintershof-West), but the sample size was limited (Figure 8). *Diaceratherium tomerdigense,* another species of teleoceratines, was also studied here, but the species is only found in one locality (Tomerdingen, MN1).

**Figure 8:**
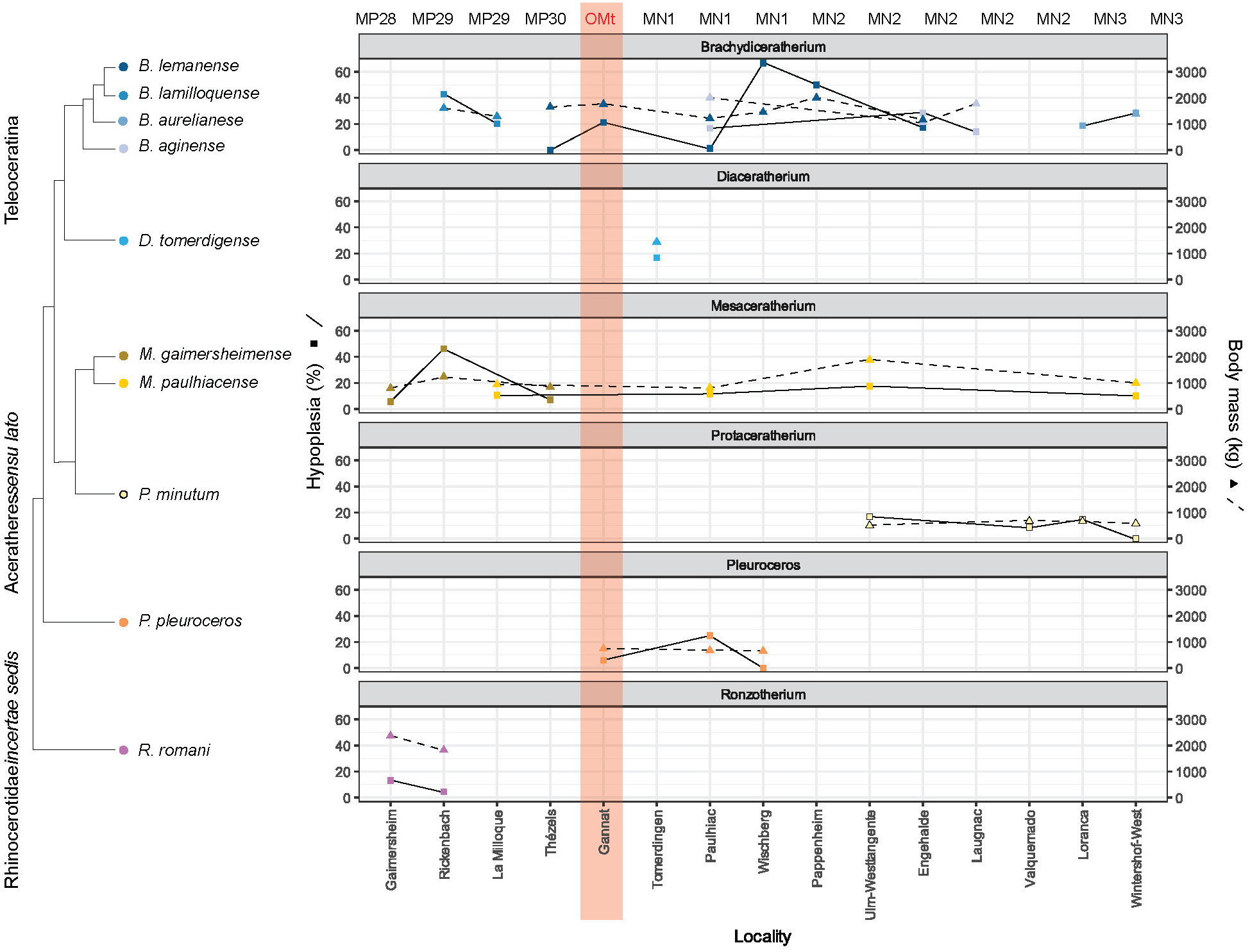
Hypoplasia prevalence (full line) and body mass (dashed line) by species and locality. Phylogenetic relationships based on the literature [99,108–110].

Regarding, the aceratheres *sensu lato*, *Mesaceratherium gaimersheimense* is only recorded during Late Oligocene. This species exhibits stable mesowear median (2 at all localities) during the interval and similar trends for body mass andhypoplasia prevalence, with a peak at Rickenbach for the first two variables (1240 kg, 46.15 %), while Gaimersheim and Thézels had similar values (about 800 kg, < 10 %). During Early Miocene, the aceratheres *sensu lato* included *M. paulhiacense* and *P. minutum*. The first one shows a peak in all parameters at Ulm-Westtangente (1880 kg, 17.45 %, 2.5), and similar values of body mass and hypoplasia prevalence at the other localities (800-1000 kg, 10-12 %; Figure 8). *Protaceratherium minutum* has similar mean body mass estimates at all four localities where it was recorded (Ulm-Westtangente, Valquemado, Loranca del Campo, and Wintershof-West), between 500 and 670 kg. Mesowear median and hypoplasia prevalence are similar for Ulm-Westtangente and Loranca del Campo (2, ∼ 15 %), but lower at Valquemado (8.51 %, 0). Hypoplasia prevalence was null at Wintershof-West but very few specimens were available (Figure 8).

Lastly, two species of basal rhinocerotidae were studied. *Pleuroceros pleuroceros* had similar mean body mass estimates (700 kg) and median mesowear ruler (2) at Gannat, Paulhiac (only mass), and Wischberg (Figure 8, Fig 3 ESM2). Hypoplasia prevalence was low or null hypoplasia except at Paulhiac (25 %), but the sample size for this species was very limited. *Ronzotherium romani* is a strictly Oligocene taxa. Body mass drastically diminished during the Late Oligocene (Gaimersheim: 2370 kg → Rickenbach: 1820 kg), similarly to hypoplasia prevalence (Gaimersheim: 13.19 % → Rickenbach: 4.55 %), but contrary to median mesowear that increased during this interval (Gaimersheim: 1.5 → Rickenbach: 4; Figure 8, Fig 3 ESM2).

## Discussion

### Dietary preferences and niche partitioning of the rhinocerotids studied

All rhinocerotids studied fell within the range of C3 feeding for stable isotopes (δ^13^C_diet_ < −22 ‰) [71], and in the range of browsers (0-2) or mixed-feeders (0.4-2.74) for mesowear [55]. Although extant grazers have reported mean mesowear scores as low as 2.09 (*Kobus ellipsiprymnus*, *Redunca redunca*), a grazing diet seems unlikely for late Oligocene – early Miocene rhinocerotids in Europe. Indeed, even though C4 grasses were present locally in Southern Europe as soon as the early Oligocene [72], they were never dominant in Europe [73–75]. Regarding the second type of grasses, C3 grasslands were also limited in Europe at that time, as most of the continent was covered by forests and woodlands [75]. Moreover, C3 grasses contain lower levels of fibers, silica, and toughness than their C4 counterparts [76], and hence C3 grazing should result in a lower abrasion load and mesowear scores.

Mesowear score is rather stable across time, space and taxa, with most species having a median mesowear score around 2 at the different localities. Similarly to other studies [77–79], we did not find a clear relationship between body mass and mesowear, suggesting once again that the classical assumption of greater tolerance for a lower quality diet in larger species lacks empirical support [80].

At some localities, we highlighted some differences in the feeding preferences of the rhinocerotid specimens associated, which could indicate a potential niche partitioning (Figure 9). On the contrary, potential competition for resources or different niche partitioning strategies was hypothesized at others. Each locality is detailed and discussed by chronological order. The dietary categorization is based on previous studies on extant herbivores (microwear: [55,56,61,65]).

**Figure 9:**
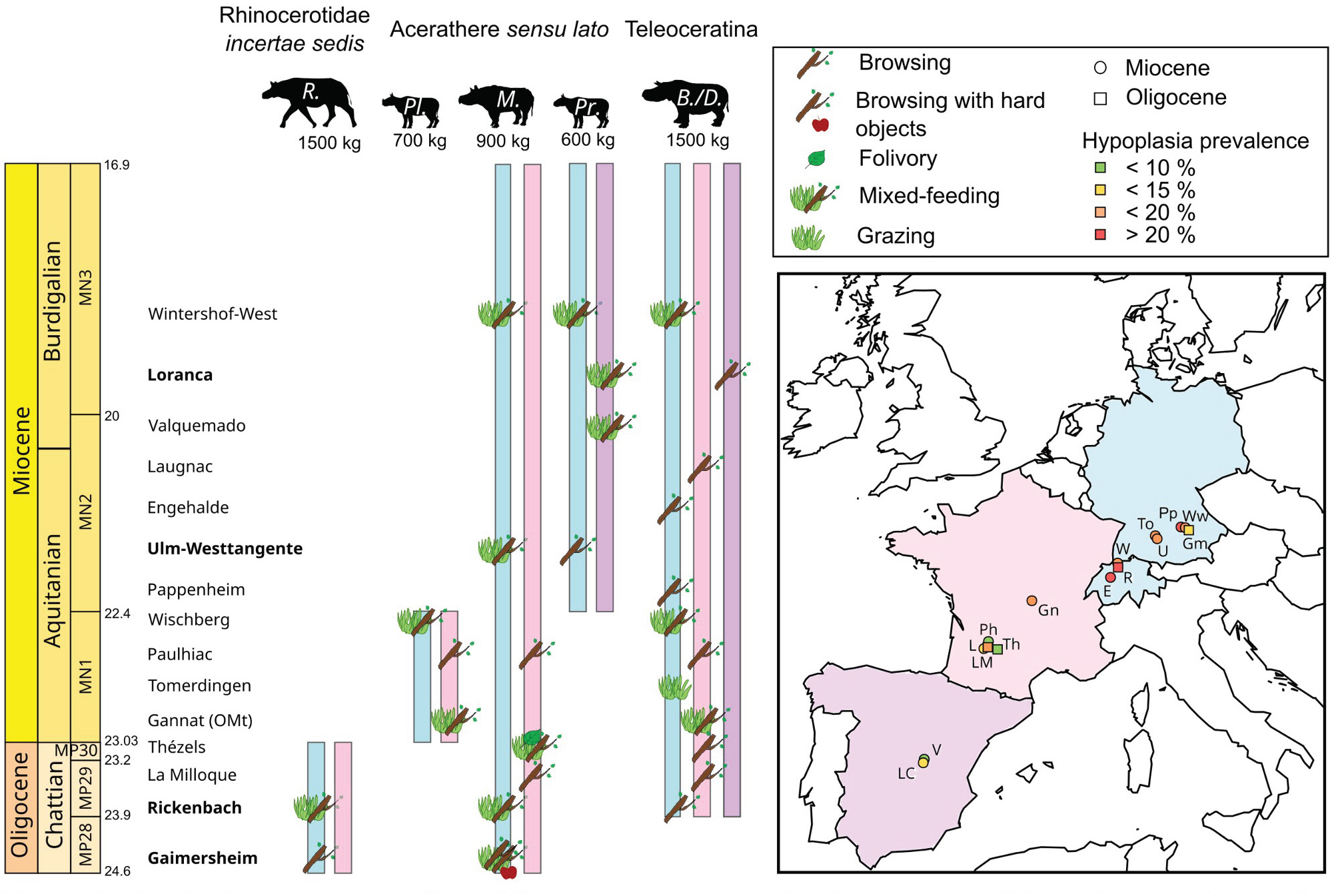
Spatio-temporal evolution of the paleobiology and paleoecology of the rhinocerotids during the Oligoce-Miocene transition. Distinct paleoecological preferences at the localities in bold. Color code by geographical provenance: blue – Central Europe (Germany and Switzerland), pink – Western Europe (France), and purple – Iberian Peninsula (Spain). <u>Abbreviations in the silhouettes:</u> *R.* - *Ronzotherium*, *Pl. -* Pleuroceros*, M*. *- Mesaceratherium*, *Pr.* - *Protaceratherium*, *B./D.* - *Brachydiceratherium* / *Diaceratherium*. <u>Abbreviations in the map:</u> E – Engehalde (MN2), Gm - Gaimersheim (MP28), Gn – Gannat (MP30-MN1), L – Laugnac (MN2, reference), LC – Loranca del Campo (MN2), LM – La Milloque (MP29), Ph – Paulhiac (MN1, reference), Pp – Pappenheim (MN1), R – Rickenbach (MP29, reference), To – Tomerdingen (MN1), Th – Thézels (MP30), U – Ulm-Westtangente (MN2), V – Valquemado (MN2), W – Wischberg (MN1), Ww – Wintershof-West (MN3).

At **Gaimersheim (MP28)**, the median mesowear of *M. gaimersheimense* (2, n = 4) and the DMTA (low Str: high anisotropy) suggest a more abrasive diet than that of *R. romani*. The carbon isotopic content of the diet was also different between both species, but only two specimens of *M. gaimersheimense* were studied. Dietary preferences were clearly separated for the two rhinocerotids at Gaimersheim, with *M. gaimersheimense* being a mixed-feeder including tougher and harder objects than the browsing *R. romani*. In a previous study [72], Heissig compared *M. gaimersheimense* to *Diceros bicornis* (extant black rhinoceros) based on tooth morphology, and proposed a similar generalist diverse and adaptable lifestyle, whereas for *R. romani* he only concluded that this species was not a steppe grazer. For the first comparison, *D. bicornis* has a similar body mass (800 – 1500 kg) [82], but a diet including more abrasive and hard objects, as suggested by mesowear scores [83] and DMT [61].

At **Rickenbach (MP29)**, both dental wear proxies suggested different feeding preferences, although not statistically significant (small sample). *Ronzotherium romani* had the highest median mesowear ruler score (4, n = 3), but low anisotropy (high Str values > 0.3). The high values of complexity and heterogeneity of the complexity suggest the inclusion of hard items and a certain variety in the diet, consistent with mixed-feeding. A previous study at Rickenbach inferred a short vegetation browsing diet for *R. romani* due to its low hypsodonty index (here HI = 1) and low head posture [84], but this would be more similar to the feeding behavior of the species at Gaimersheim.

*Brachydiceratherium lamilloquense* had the lowest median mesowear score (2, n = 2) consistent with the DMTA signature (high Str, medium Asfc and Hasfc) and pointing towards browsing habits. The low hypsodonty index (here 0.89, brachyodont) and the intermediate head posture were used in a previous study to infer high level browsing [84]. Eventually, the third species *M. gaimersheimense*, recognized in a recent revision [85], had a moderate median mesowear score (2, n = 3), high complexity (Asfc > 2) and low anisotropy (Str > 0.3) suggesting browsing. [38][86–88][81][11,84]

At **La Milloque (MP29)**, both rhinocerotid species overlap for carbon isotopic content and DMT, although some slight differences can be noted. The mesowear median however highlights a higher abrasivity in the diet of *M. paulhiacense* (2, n = 7) compared to *B. lamilloquense* (1, n = 5). Both species appear to have been browsers, but probably fed on different plants or plant parts (different body mass, feeding height and mesowear). The study of the Moschidae species at La Milloque revealed very diverse dietary preferences, with the largest species in the range of extant grazers and smallest probably folivore [89]. These preferences are rather distinct from the ones infer for the rhinocerotids here, which supports the conclusion of the authors of a relatively heterogeneous environment at the locality.

At **Thézels (MP30)**, the mesowear and isotopic content are only available for *M. gaimersheimense* and indicate browsing or mixed feeeding. The DMTA however, points towards a tough, abrasive diet (Str < 0.2). This pattern (low mesowear score, high anisotropy) has previously been linked to folivory [34,61], but could also be the result of C3 grazing or seasonal variations in the diet. C3 grazing has already been hypothesized for some brachyodont Moschidae at La Milloque based on their microwear [89]. Seasonality has previously been discussed at Thézels [90] and our results concur to a great aridity at this locality (see next section).

Only one specimen of *B. lemanense* was examined for microwear and falls in the range of variation of *M. gaimersheimense*, suggesting at least a partial overlap in the dietary preferences of both species. Similar paleoecologies have already been proposed for both rhinocerotids at Thézels: Blanchon et al. [91] interpreted them as browsers and cursorial forest dwellers. However, the body mass of these rhinocerotids are very different (850 kg for *M. gaimersheimense* and 1650 kg for *B. lemanense*), which could point to another strategy for niche partitioning.

At **Gannat (MP30 to MN1)**, dental wear (meso- and micro-wear) was studied for *B. lemanense* and *P. pleuroceros*, while isotopic content could only be retrived for the first one. The median mesowear scores of both species are equal to 2, suggesting a potential overlap of the feeding preferences in the browser to mixed-feeder range. Only one specimen of *P. pleuroceros* could be studied for DMTA, and its signature is within the range of *B. lemanense* microwear, confirming the potential dietary overlap of both species.

Rhinocerotid material from the earliest Miocene (MN1) was scarce. At **Tomerdingen (MN1)**, only one species (*D. tomerdingense*) is recorded, for which only mesowear scoring was possible. The only molar examined had very high score suggesting grazing. At **Paulhiac (MN1)**, the median mesowearsuggested soft browsing for both *Brachydiceratherium* species and the DMTA highlighted partial overlap between *B. lemanense* and all other species, notably for values of complexity. At **Wischberg (MN1)**, very few specimens could be analyzed and do not allow for species comparison (only *Brachydiceratherium* for DMTA and only *P. pleuroceros* for mesowear), but both species are in the mixed-feeders range.

At **Pappenheim (MN2)**, there is only one species (*B. lemanense*), for which mesowear (2, n = 1) and DMTA (high Str, high Asfc, low HAsfc) suggest browsing. At **Ulm-Westtangente (MN2)**, there was a clear partitioning in the diet and/or habitat as highlighted by carbon isotopes and mesowear: *M. paulhiacense* had a more abrasive diet and was probably a mixed feeder, while *P. minutum* was a browser. The DMTA revealed less differences as discussed by Hullot et al. [92]. At **Engehalde (MN2)**, very few specimens of rhinocerotids were found, limiting the analyzes. The comparison between species is not possible, as the DMTA only include one specimen of *B. aginense* (grinding facet only) and the mesowear was assessed only for *B. lemanense*. Both species are very similar and were probably browsers, in a humid wooded habitat close to steady rivers and swamp area [93]. At **Laugnac (MN2)**, only one species was studied (*B. aginense*), for which low mesowear and DMT signature (high Str, moderate Asfc, low HAsfc) point toward soft browsing. At **Valquemado (MN2)**, only one species is found (*P. minutum*). The isotopes and DMTA suggest mixed-feeding habits, contrasting with the very low mesowear estimated on one specimen (0, n = 1).

At **Loranca del Campo (MN3)**, the mesowear and isotopic analysis only include the more abundant species: *P. minutum*. The DMT reveals different dietary preferences for both species (shearing facet): *P. minutum* has a higher anisotropy and slightly higher complexity, highlighting a tougher diet (mixed-feeder) than for *B.* aff. a*urelianense* (browser). The DMT of *P. minutum* between Valquemado and Loranca del Campo denotes a higher anisotropy at the latter (Figure 7), consistent with morphological differences (size, gracility) and the more arid conditions previously inferred at Loranca [17]. These inferences are however not supported by our results, suggesting similar body mass (Table 3) and similar environmental conditions at both localities (see next section). Eventually, at **Wintershof-West (MN3)**, the DMT of all three species overlaps in the mixed-feeding range. The mesowear of *B. aurelianense* (3, n = 1) and *M. paulhiacense* (1, n = 1) are however very distinct but the sample size is restricted.

### Paleoenvironmental conditions

The analyses of the isotopic content (carbon and oxygen) in the carbonates of the rhinocerotids’s enamel allow for some paleoenvironmental insights (Table 4). However, these results are only partial, as the sample is limited to rhinocerotids at some localities, and should be completed by results on other taxa to provide more robust reconstructions. There was a clear shift towards higher values of δ^13^C_diet_ for rhinocerotoids between La Milloque (MP29) and Thézels (MP30; Figure 6). This suggests the establishment of more open and arid conditions during the latest Oligocene and persisting into the Early Miocene. Mean annual temperatures (MAT) were rather warm at all localities but Gaimersheim, for which MAT was between 1.4 and 6°C lower than that at other localities (Table 4). Regarding precipitation, the mean annual precipitation (MAP) suggest rather arid conditions at Thézels, Gannat, Valquemado, and Loranca del Campo. The values estimated with Kohn’s equation [42] are extremely low at these localities (< 100 mm), and to a lesser extent for the whole dataset (< 500 mm). This could be explained by the consumption of certain plants (for instance C4) [42], as well as the inclusion of altitude in this equation, which is difficult to estimate for fossil sites. Results for MAP are less drastic with Rey’s equation [43], although the equation was built using the same dataset from Kohn [42]. In all cases, the paleoenvironmental conditions seem relatively favorable at all localities investigated, consistent with the low (10 %) to moderate (< 20 %) prevalence of hypoplasia for most rhinocerotids and localities (Table 3; Figure 5).

**Table 4:** Reconstruction of the mean annual temperature (MAT) and precipitation (MAP) based on the isotopic content of the carbonates of the rhinocerotids’ enamel.

|  | Datation | MAT – Tütken et al. [47] (°C) | MAT – Skrzypek et al. [48] (°C) | MAP – Kohn [42] (without neg. values; mm/year) | MAP – Rey et al. [43] (mm/year) |
| --- | --- | --- | --- | --- | --- |
| **Gaimersheim** | MP28 | 13.6 | 12 | 273 | 857 |
| **Rickenbach** | MP29 | 17.4 | 14.5 | 282 | 896 |
| **La Milloque** | MP29 | 19.2 | 15.8 | 364 | 973 |
| **Thézels** | MP30 | 19.8 | 16 | 95 | 460 |
| **Gannat** | MP30-MN1 | 19.7 | 16.1 | 62 | 586 |
| **Ulm-Westtangente** | MN2 | 15.6 | 13.4 | 317 | 637 |
| **Valquemado** | MN2 | 20.6 | 16.5 | 72 | 513 |
| **Loranca del Campo** | MN3 | 20.4 | 16.4 | 51 | 530 |

Our paleoenviromental insights can be confronted to the global climate predictions for the Late Oligocene – Early Miocene. The Late Oligocene Warming occurred between the MP26 and MP28 (∼26.5 to 24 Ma) [3], and was responsible for a temperature increase in terrestrial environments of up to 10°C [4,90,91]. Localities from our dataset all postdate this event, except maybe Gaimersheim (MP28), but the estimated MAT and MAP at this site suggest temperate conditions (Table 4). Interestingly, localities from the MP29 (Rickenbach, La Milloque) and MP30 (Thézels) have warm MATs (> 15°C), suggesting the persistence of warm conditions during the latest Oligocene, at least locally. However, seasonal arid conditions have been hypothesized at Rickenbach, due to the presence of evaporite levels [38]. Interestingly, Rickenbach has one of the highest hypoplasia prevalence of the dataset (13/49; 26.53 %), and notably *B. lamilloquense* (6/14; 42.86 %) and *M. gaimersheimense* (6/13; 46.15 %). Both teleoceratines and aceratheres *sensu lato* having been inferred as aquaphiles [86–88], which might explain the higher stress levels observed for these species compared to *R. romani* (1/21; 4.55 %), supposedly more adapted to arid conditions [81]. Moreover, the Rickenbach level (reference of MP29) corresponds to the beginning of the “Terminal Oligocene Crisis” [18] and a faunal turnover [11,84], which might have created stressful conditions for the rhinocerotids, notably through competition.

Moreover, he MAP estimates show a drastic drop between La Milloque and Thézels (Table 4), indicating an increase in aridity during the MP30. Interestingly, the sediments at Thézels (tertiary limestone) and in the region (Quercy) suggest a seasonal aridity with periodic flooding [90]. Surprinsignly, the hypoplasia prevalence at Thézels was one of the lowest in our dataset (11/171; 6.43 %), although such conditions of periodic flooding have been linked with high hypoplasia prevalence in rhinocerotids at other sites [34,92].

The MP30 has been correlated to the interval between 23.2-23.03 Ma [5,6], which coincides with the drop in δ^18^O_benthic foraminifera_ documenting the Mi-1 event [95]. Hence, this change in isotopic content could document the beginning of the Mi-1 glaciation event and associated vegetation changes. A few individuals from these two localities have been serially sampled to investigate seasonality. Most specimens display a sinusoidal variation of both δ^13^C_CO3_ and δ^18^O_CO3_ (ESM2 Figs 7 and 8), which could attest of some seasonality. Intra-individual variations observed for δ^18^O_CO3_ (0.6 to 2.5 ‰, mostly around 1 ‰) were greater than that of δ^13^C_CO3_ (0.2 to 2.4 ‰ but mostly around 0.5 ‰). Some teeth sampled (first and second molar, fourth premolars) might have recorded some weaning signal, but their variations are similar to that of third molars. We tested the correlation between the δ^13^C and δ^18^O values in the enamel carbonates, which would indicate a consistent influence of seasonal precipitation on the plants consumed. We only found one at La Milloque (rho = 0.5698249, S = 13273, p-value = 3.72e-06), most probably as more specimens were studied and with more samples per tooth.

Regarding Gannat, many discussions are found in the literature about dating problematics [96–98]. The presence of *R. romani* and *Eggysodon pomeli*, typical Oligocene taxa [99], would suggest an Oligocene age, whereas the other two species studied here (*P. pleuroceros* and *B. lemanense*) indicate early Miocene age. However, the specimens of *R. romani* and *E. pomeli* come from a different pit that is probably not contemporaneous. The isotopic content of the enamel of *B. lemanense* suggest rather warm and arid conditions, that would be consistent with the installation of the warmer and more humid conditions of the Aquitanian, shortly after the Mi-1 event [2,7]. However, the conditions at Gannat are also very similar to that of Thézels, which could advocate for a latest Oligocene age. Hence, absolute dating appears crucial at Gannat, to untangle the situation. Two specimens were also serially sampled, one to detect a potential weaning signal (SpA) and one to investigate seasonality (SpB). Variation for δ^18^O_CO3_ were greater than that of δ^13^C_CO3_ in both specimens, similar to the results at La Milloque and Thézels (ESM2 Figs 4 to 6). Variations in the values of δ^18^O_CO3_ in SpB are abrupt and might indicate a strong seasonality, although they are in the range of the weaning shift observed for SpA (ESM2 Fig 6).

In addition to the documentation of chronological aspects, the δ^18^O values and estimated MATs might document geographical variations within Europe. Indeed, if localities are ordered spatially (Gaimersheim, Ulm-Westtangente, Rickenbach, Gannat, Thézels, La Milloque, Valquemado, Loranca del Campo; see Figure 1, 4, and ESM2 Fig 7), the MATs are increasing from North-East to South-West. This separates localities from Central (Germany, Switzerland) and Western Europe (France, Spain), with warmer and relatively drier conditions in the latter (except for La Milloque). Indeed, both localities of the Iberian peninsula, Valquemado and Loranca del Campo, had high MATs and low MAPs (4), consistent with previous findings indicating that this region was warmer and more arid than the rest of Europe already during the Early Miocene [75,100]. Interestingly, the DMTA also support this, as specimens of *P. minutum* from both Iberian localities have slightly higher complexity and lower anisotropy (on the grinding facet) than at Ulm-Westtangente or Wischberg (Figure 7). Regarding hypoplasia prevalence, rhinocerotids from Central European localities seem more affected, which does not reflect this potential aridity gradient.

The results of this study highlight several trends in the evolution of body mass, dietary preferences and susceptibility to stresses during the Oligocene-Miocene transition. Rhinocerotids in our dataset show a tendency of decreasing body mass during Late Oligocene, that is prolonged to the MN1 for some species (Figure 8). During the Early Miocene, all trends (increase, stability, decrease) are observed depending on the species, contrary to what we expect (increase in body mass during the Early Miocene along with the appearance of mediportal forms). Interestingly, hypoplasia prevalence and body mass co-variated, suggesting a higher stress susceptibility in bigger species. This result is not surprising, as bigger species are more vulnerable to environmental changes, as they have less babies, a longer gestation time, and they more sensitive to habitat and population fragmentation [101,102]. However, other parameters might be responsible for this correlation, like a bigger size at localities with harsher conditions (Bergmann’s rule: bigger species or population in colder climates [98]). This result however challenges the typical assumption that bigger species can buffer seasonality changes and tolerate lower nutritious diet [15], and show that these stressful conditions also impact larger species.

The prevalence of hypoplasia is indeed greater around the Oligocene-Miocene limit (Gannat, Wischberg, Paulhiac) compared to Late Oligocene and Early Miocene where it oscillates between 5-15 % for most species at most localities (Table 3). This overall low to moderate prevalence is in line with our paleoenvironmental insights (Table 4), as well as with the warm and humid during the Aquitanian initiating the conditions of the Miocene Climatic Optimum [2]. Interestingly, Teleoceratine (here *Brachydiceratherium* and *Diaceratherium*) were the most affected taxa, recalling the pattern observed for *Brachypotherium* during the Early-Middle Miocene [104]. Teleoceratine have often been associated with humid environments and conditions [87,88], although a semi-aquatic lifestyle is not supported [105,106]. This water dependency might make these taxa more vulnerable to seasonality and aridity. Sadly, the direct comparision with localities of the MN4 onwards is difficult due to an important turnover in rhinocerotid species following the Proboscidean Datum Event [107], and marking the end of the Oligocene-Miocene transition [11].

## Conclusions

The results of this study highlight several trends in the evolution of body mass, dietary preferences and susceptibility to stresses during the Oligocene-Miocene transition. Changes recorded in the enamel of rhinocerotids during this interval, suggest seasonal aridity at several localities (Thézels, Gannat, Valquemado, and Loranca del Campo) and changes in the vegetation during the latest Oligocene. An increase in abrasivity is also observed in the DMT of several species during this interval, suggesting a shift in dietary preferences and/or habitats during the lastest Oligocene. The reconstructed paleoenvironmental insights indicate favorable conditions, with warm mean annual temperatures (> 15 °C) at most localities studied. This is consistent with the low to moderate prevalence of hypoplasia (5-20 %). Even if all rhinocerotids were C3 feeders, with browsing to mixed-feeding preferences, we highlighted differences in dietary and or habitat preferences that could indicate niche partitioning at some localities (Gaimersheim, Ulm-Westtangente, and Rickenbach). Our study showed great changes in the paleoecology of the rhinocerotids during the Oligocene-Miocene transition, in line with taxonomic and morphological changes previously noted, and with the global climatic and paleoenvironmental changes at that time.

## Supporting information

ESM2

ESM1

## Acknowledgements

The present work is part of a post doctoral project funded by a Research Fellowship of the Alexander von Humboldt Foundation (Germany). Two SYNTHESYS grants have also been obtained (FR-TAF_Call4_057, ES-TAF-TA4-781) and funded partly this work.

We are indebted to the curators and collection managers of the institutions that we visited: Eli Amson (SMNS Stuttgart), Susana Fraile (MNCN Madrid), Christine Argot and Guillaume Billet (MNHN Paris), Jérome Surrault (University of Poitiers), Christophe Borrely (CECM Marseille), Ursula Menkveld (NHM Bern), Emmanuel Robert (University of Lyon), Didier Berthet and François Vigouroux (Musée des Confluences de Lyon), Loic Costeur and Florian Dammeyer (NHMB Basel), Jean-Marc Pouillon (Rhinopolis Gannat) and Florence Lamera (Paleopolis Gannat), Pia Geiger-Schütz and Peter Flückiger (Naturmuseum Ölten, Haus der Museen).

## Notes

### Competing Interest Statement

The authors have declared no competing interest.

### Summary of Updates

The R. romani specimen from Gannat was revised to Egyssodon pommeli and as such removed from the study because it is not a Rhinocerotidae (but a Rhinocerotoidea). According to reviewers comments, we modified the previous version notably: - including some figures from the supplementary into the text; - using median for mesowear as it is ordinal categorical and the use of mean does not make sense mathematically; - adding clearer work hypotheses in the introduction; - explaining the context better in the introduction.

