## Supplementary material for "Evolutionary paleoecology of European rhinocerotids across the Oligocene-Miocene transition": ESM2

**Supplementary data of “Evolution of the paleoecology of European rhinocerotids across the Oligocene-Miocene transition”**

Details on localities (chronological order) and species studied

**Gaimersheim** refers to a fossil locality originating from the filling of a system of karst fissures in the dolomite of the upper Weißjuras. It was found in the Gaimersheim quarry (48.820145° N, 11.379961° E) near Ingolstadt in Bavaria (southeastern Germany). It has been dated to the Chattian (late Oligocene) [1] and the fauna has been correlated to the Mammal Paleogene (MP) reference level 28. Two species of rhinoceros have been recognized: *Ronzotherium romani* and *Mesaceratherium gaimersheimense*. The rhinoceros material consists in 179 teeth and tooth fragments and 15 jaw pieces, excavated between 1944 and 1954, and stored at the Bayerische Staatssammlung für Paläontologie und Geologie München.

**Rickenbach** was discovered in 1897 during the construction of a draw well. It is found in the sandstones of the Rickenbach pit, which is also known locally as the "Huppergrube", west from Olten and at the northern boundary of the foreland molasse in Switzerland [2,3]. Rickenbach is the type locality for the MP29 dated to around 23.5 Ma. The Cricetidae suggest a slightly older age for Rickenbach than for La Milloque. Three species of rhinoceros have been identified in the latest revision of the Olten material [4]: *Ronzotherium romani*, *Mesaceratherium gaimersheimense*, and *Brachydiceratherium lamilloquense*.

**La Milloque** is a fossil locality from southwestern France discovered in 1868. It is characterized by a set of molasses deposits called ‘Molasses de l’Agenais’, which underlie the Lower Miocene white limestone called ‘Calcaire blanc de l’Agenais’ [5,6]. It has yielded an abundant vertebrate fauna correlated to the MP29 and dated to about 24 Ma [7]. Two species of rhinocerotids are found at La Milloque: *Mesaceratherium paulhiacense* and *Brachydiceratherium lamilloquense.*

**Thézels** is located south of Cahors on the north-eastern margin of the Aquitaine basin (southwestern France; [8]. The fauna is correlated to the MP30. Two species of rhinocerotids are found at Thézels: *Mesaceratherium gaimersheimense* and *Brachydiceratherium* aff. l*emanense [9]*.

**Gannat** is composed of several fossiliferous pits found 20 km West from Vichy (Central France), in the northern part of the Massif Central. Gannat quarries are located at the top of the hills “Puy de Clermont” and “Mont Libre” Southwest of the city of Gannat [10,11]. The vertebrate faunas from the different pits are correlated to the late Oligocene (“Gannat sommet”, Chattian, MP30) and to the early Miocene (“Gannat supérieur”, Aquitanian, MN1) [12]. The rhinoceros assemblage is dominated by *Brachydiceratherium lemanense* but a few specimens of *Pleuroceros pleuroceros* are also found. Two other typically Oligocene rhinocerotoids are also reported at Gannat *Ronzotherium romani* and *Eggysodon pomeli*, but from different pits [10].

**Tomerdingen** is a fissure filling, located 11 km North-West of Ulm (Baden-Württemberg, Germany) in the Tomerdingen quarry. The fauna has been correlated to the MN1 [13] and has yielded one species of rhinocerotid: *Diaceratherium tomerdingense* [14].

**Paulhiac** is a fossil site from southwestern France (Lot-et-Garonne), located about 6 km north-east to the village of Montflanquin [15]. This locality is the reference for the MN1 [5,16]. The rhinocerotid material has been attributed to four species: *Brachydiceratherium aginense*, *Brachydiceratherium lemanense*, *Mesaceratherium paulhiacense*, and *Pleuroceros pleuroceros* [17,18].

**Wischberg** is a Swiss locality situated in the Bern Canton (latitude 47.199157894°/longitude 7.763943664°) and found in the Lower Freshwater Molasse [19]. It has yielded one of the richest fauna in Switzerland of the early Miocene, and it has been correlated to the MN1. Two species of rhinocerotids are found in association at this site according to the recent revision of the material [18]: *Brachydiceratherium lemanense* and *Pleuroceros pleuroceros*.

**Pappenheim** is fissure filling locality of Southeast Germany. The faunal assemblage has been correlated to the MN2. One species of rhinoceros, initially identified as *Pleuroceros pleuroceros* [17], but is here referred to *Brachydiceratherium lemanense* based on size and dental characters.

**Ulm-Westtangente** is one of the richest fossil mammal localities of the early Miocene (Aquitanian: 23.04 – 20.44 Mya) [20] in Germany and even in Europe [21,22]. This fossil site was found during the construction of a road, in the Lower Freshwater Molasse sediments of the Baden-Württemberg Basin in Southwestern Germany, about 5 km North-West of Ulm (590 m above sea level, coordinates: 48.418321 N, 9.933701 E –

converted from the Gauss Kruger coordinates in original publication [21]: r 35 69 188, h 53 64 925). It has been dated to the early Miocene by correlation with the Mammal Neogene-Zone 2a (MN2a) [23,24]. Two species of rhinocerotids are recognized, although a revision is needed and there might be a third species (*Plesiaceratherium platydon*): *Mesaceratherium paulhiacense* and *Protaceratherium minutum* [22].

**Engehalde** is a Swiss fossil locality discovered in 1850 in the Bern Canton [25]. The site is in the Lower Freshwater Molasse and dated to the latest Aquitanian age, correlated to the MN2b [26]. A second contemporaneous site has been discovered recently (2006-2008) during the construction of the Neufeld tunnel in Bern city, but has yielded mostly ruminants [27]. In the historical level, two rhinocerotid species were found: *Brachydiceratherium lemanense* and *B. aginense*. The paleoreconstruction suggests a humid wooded habitat close to steady rivers and swamp areas for the historical level and a more distal well-drained floodplain, partly wooded, with open high-grassland and seasonal floods for the Neufeld level [26].

**Laugnac** is found 9 km North of Agen (Southwestern, France). It is the type locality of the MN2b [28]. Several species have been reported at this site (*Brachydiceratherium lemanense, Pleuroceros pleuroceros*, *Protaceratherium minutum*, *Plesiaceratherium aquitanicum*, *Mesaceratherium paulhiacense*), but only *Brachydiceratherium aginense* was documented by the dental material studied here [17].

**Valquemado** is a fossil site of the Cuenca province in Spain. The mammal assemblage suggests a correspondance with the MN2b. Only one species of rhinocerotid is found at this site: *Protaceratherium minutum* [29].

**Loranca** is a fossil site of the Cuenca province in Spain. The mammal assemblage suggests a correspondance with the MN3a. Two species of rhinocerotids are documented: *Brachydiceratherium* cf. *aurelianense* and *Protaceratherium minutum* [29]. Slightly more arid conditions are inferred at Loranca, compared to Valquemado.

**Wintershof- West** is a MN3 fossil locality of the North Alpine foreland in Southern Germany, close to Eichstätt (60 km North of Munich, 100 km West of the Czech border; [30,31]. Three species of rhinocerotids have been identified: *Brachydiceratherium* cf. *aurelianense*, *Mesaceratherium paulhiacense*, and *Protaceratherium minutum*. Paleoecological reconstrutions suggest a mean annual temperature above 17°C and precipitation around 858 mm. A certain forest cover is hypothesized, with slowly flowing waters and water reservoirs nearby [31].

Supplementary figures regarding results

See next page.

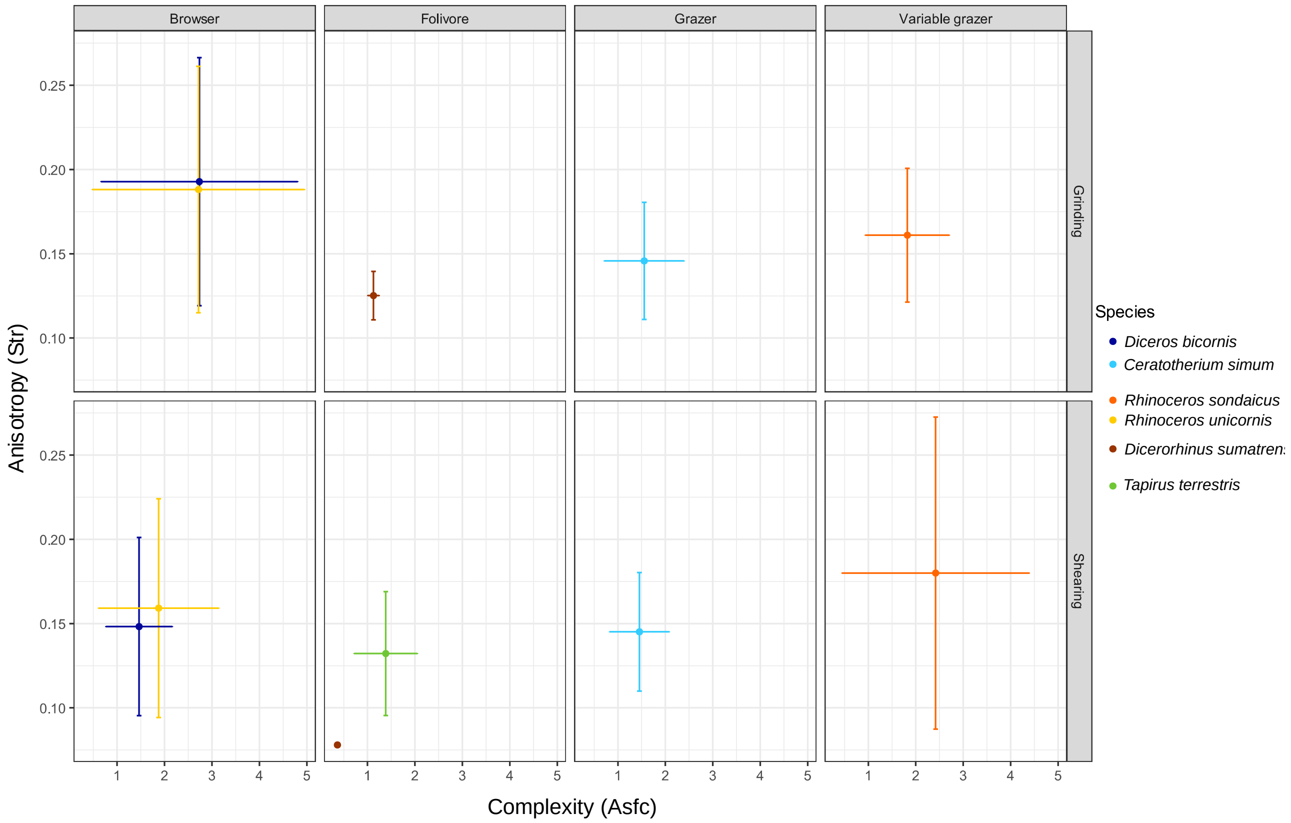


**Figure 1: DMTA of extant species.**

Dataset from Hullot et al [34] completed and reanalyzed.

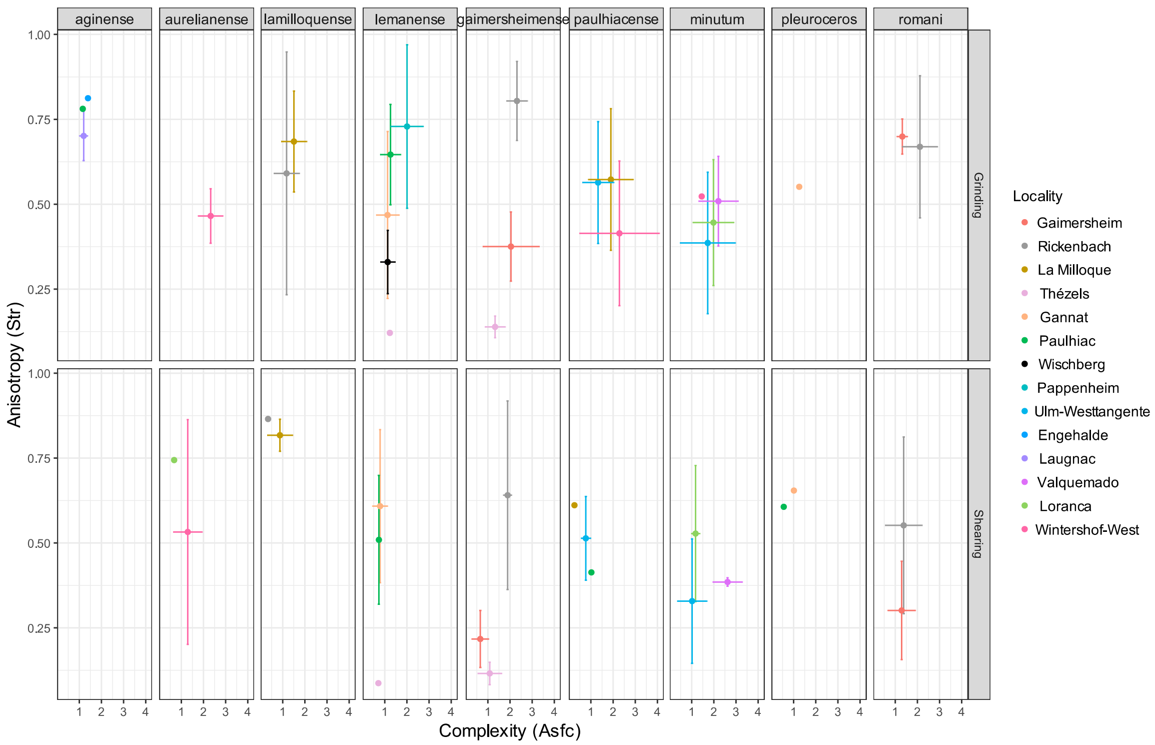


**Figure 2: Microwear (Str vs. Asfc) by species and locality.**

Phylogenetic relationships based on the literature [18,35–37].

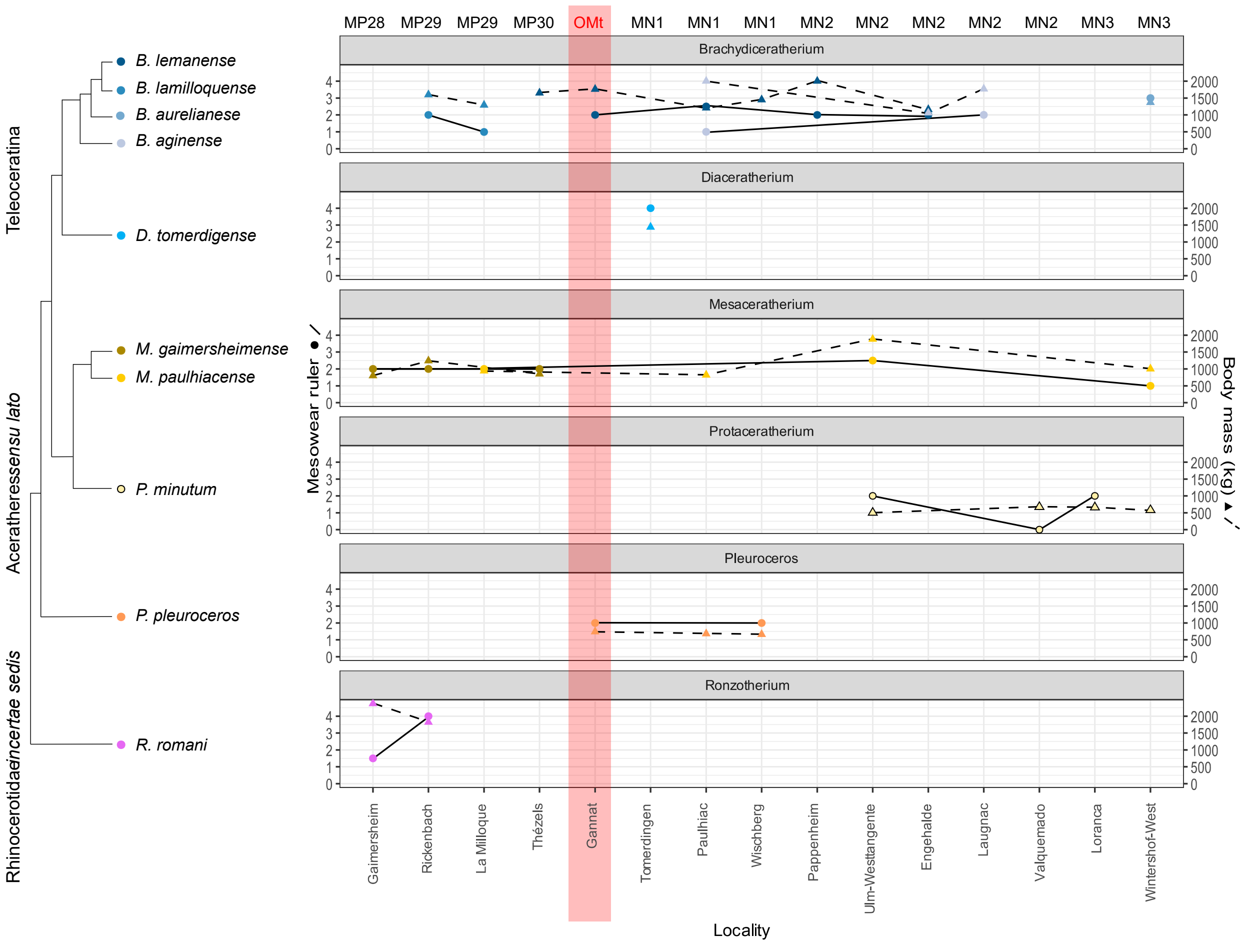

**Figure 3: Mesowear ruler and body mass by species and locality**.
Phylogenetic relationships based on the literature [18,35–37].

Supplementary figures regarding discussion: serial sampling for stable isotopic analyses

Some specimens from La Milloque (MP29), Thézels (MP30), and Gannat (MP30-MN1) were serially sampled for isotopic analyses to investigate seasonality. The results for these analyses are detailed thereafter by locality. Sinusoidal patterns are visible for many specimens. For some, only a partial sinusoid is observed, as some tooth fragments were too small to get enough samples for a sinusoidal pattern to show. For such specimens, variations of $\delta^{13}C_{CO3}$ and $\delta^{18}O_{CO3}$ values are still noted. The co-variation of $\delta^{13}C_{CO3}$ and $\delta^{18}O_{CO3}$ values was tested, as it would indicate a consistent influence of seasonal precipitation on the plants consumed.

**La Milloque:** Profiles were sinusoidal or partly sinusoidal (if not enough samples) for all specimens indicating some variations for both $\delta^{13}C_{CO3}$ and $\delta^{18}O_{CO3}$. Both parameters were correlated (rho = 0.5698249, S = 13273, p-value = 3.72e-06), indicating seasonal variations of the diet.

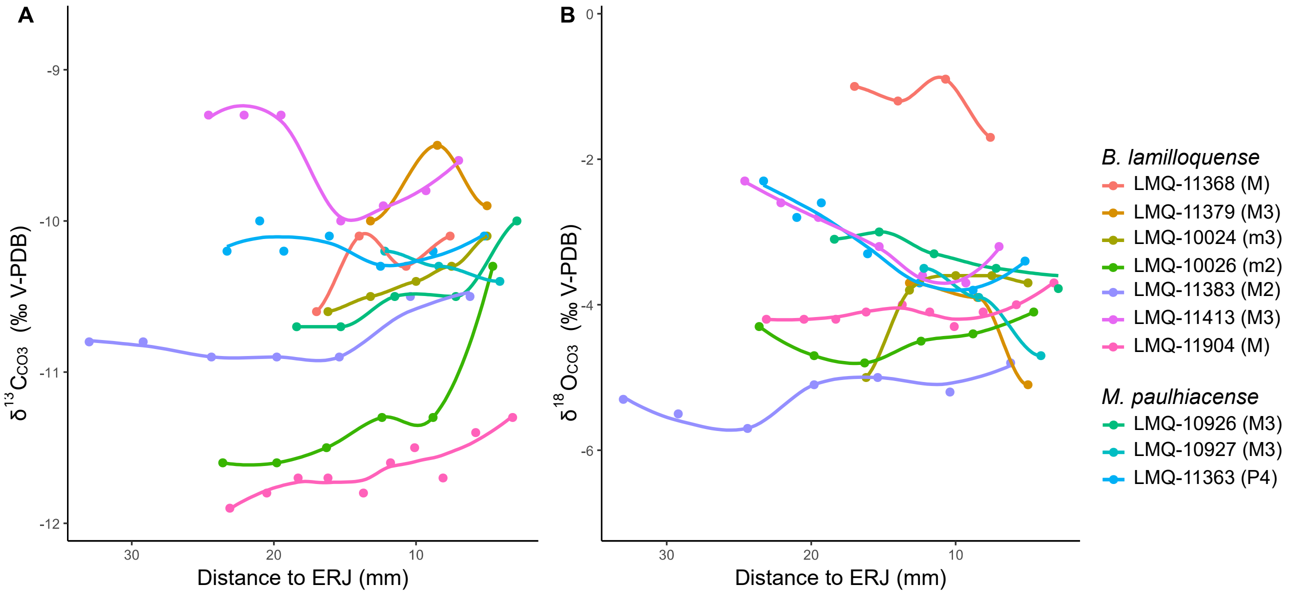


**Figure 4: Serial sampling for carbon (A) and oxygen (B) isotopes content of rhinocerotids' enamel at La Milloque (MP29).**

**Thézels:** Profiles were sinusoidal or partly sinusoidal for all specimens indicating some variations for both $\delta^{13}C_{CO3}$ and $\delta^{18}O_{CO3}$. There was no correlation (rho = -0.1633821, S = 1547.3, p-value = 0.4913) between both parameters, but it might be due to a lower sample size (both specimens and samples) than at La Milloque. The isotopic content of the rhinocerotids' enamel confirm the seasonal aridity already inferred at Thézels [8].

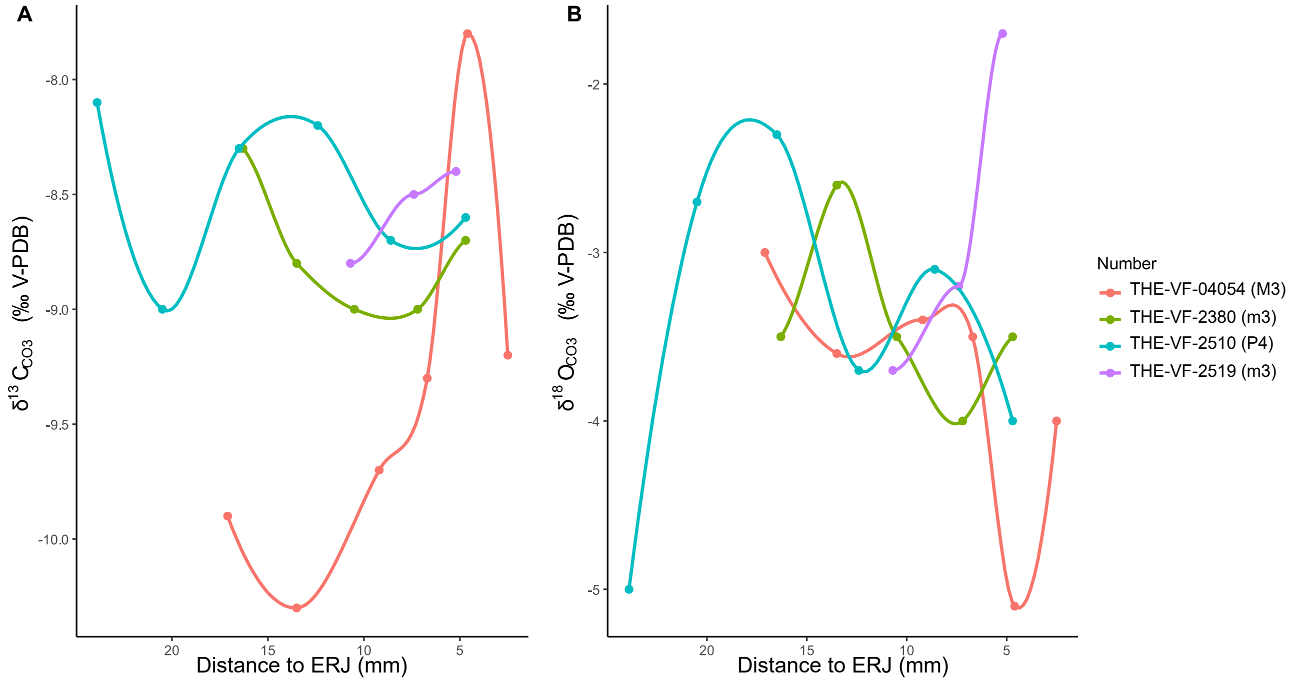


**Figure 5: Serial sampling for carbon (A) and oxygen (B) isotopes content of rhinocerotids' enamel at Thézels (MP30).**

**Gannat:** Only two specimens of *B. lemanense* were serially sampled at Gannat. This probably documents the earliest Miocene (MN1) conditions. The first specimen (SpA) only had two samples to investigate weaning signal (one before weaning and one after). For SpA we found no variations for $\delta^{13}C_{CO3}$ but a drop in values for $\delta^{18}O_{CO3}$. The second specimen (SpB) was studied to investigate seasonal variations. This revealed great variations for both $\delta^{13}C_{CO3}$ and $\delta^{18}O_{CO3}$, but without clear correlation (rho = -0.2307692, S = 147.69, p-value = 0.5502). The variations are however in a similar range to that of the weaning signal.

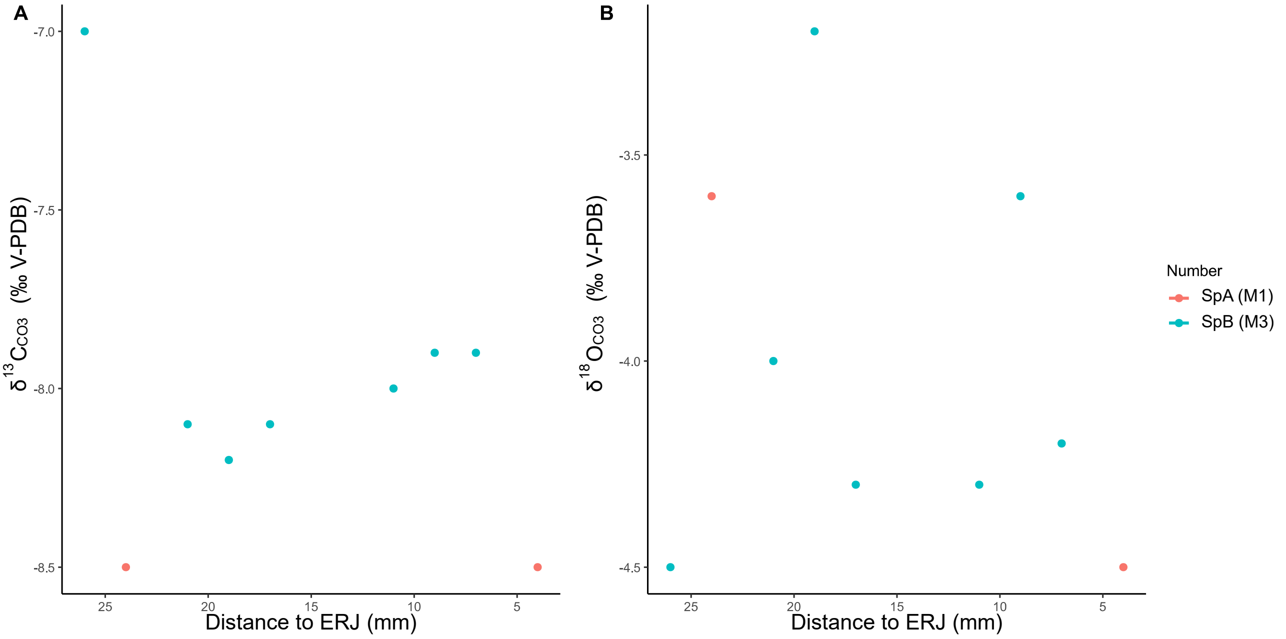


**Figure 6: Serial sampling for carbon (A) and oxygen (B) isotopes content of rhinocerotids' enamel at Gannat (MP30-MN1).**

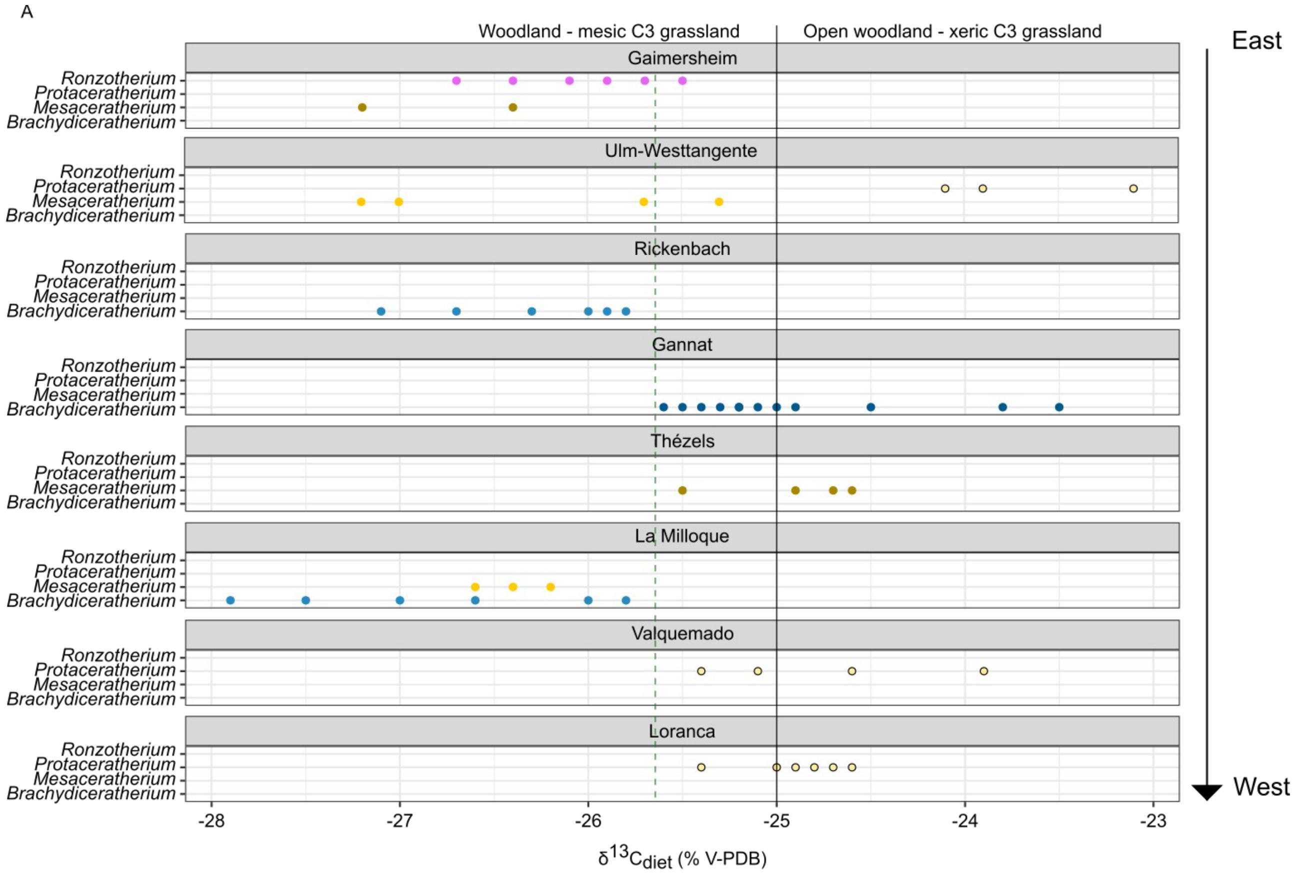

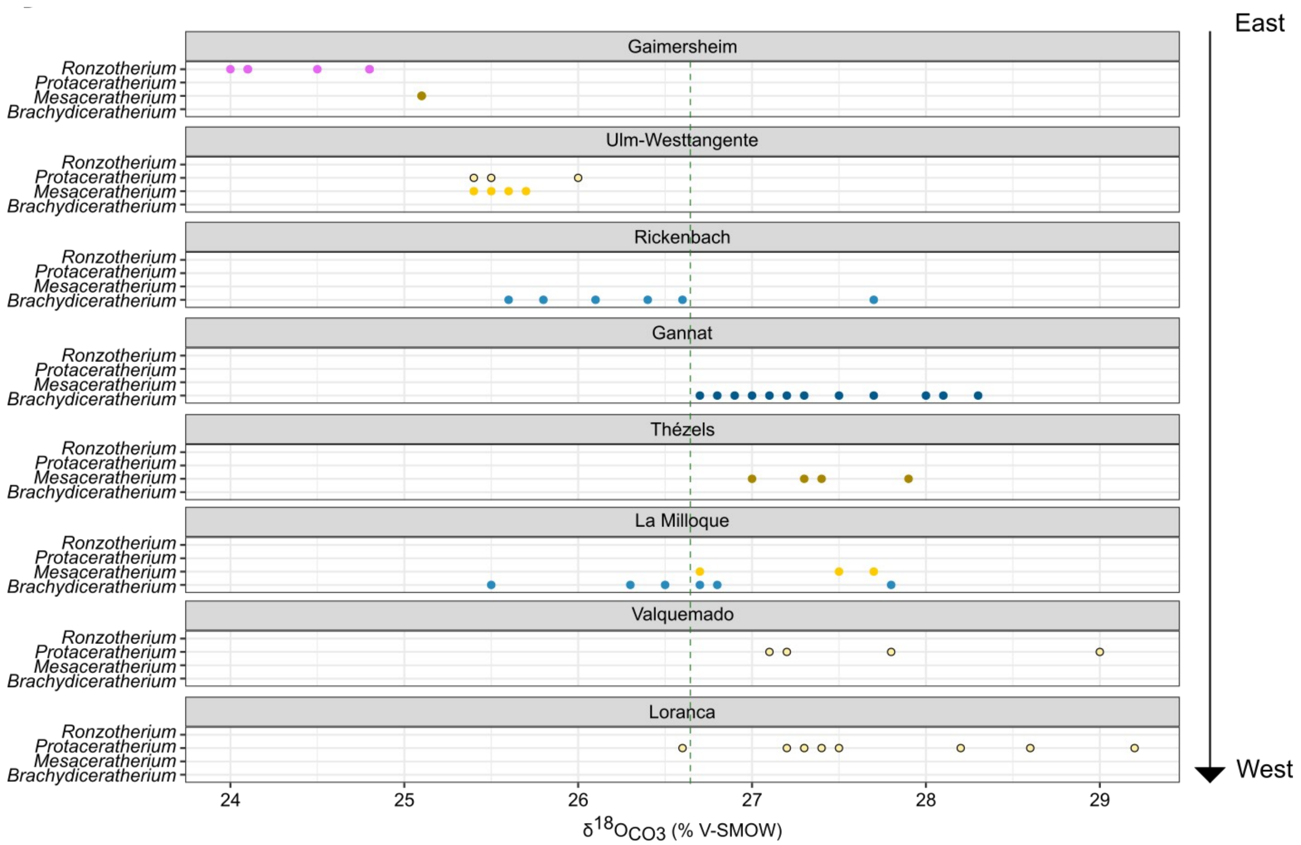


**Figure 7: Carbon (A) and oxygen (B) content of the diet and enamel of rhinocerotids from various Late Oligocene to Early Miocene localities.**
Green dashed lines: shift, black lines: cutoff between different biomes [38]. Color code by species as detailed in the figure.

*Land Mammals of Europe* (eds GE Rössner, K Heissig), pp. 9–24. München, Germany: Verlag Dr. Friedrich Pfeil.

25. Studer B. 1853 *Geologie der Schweiz. Zweiter Band. Nördliche Nebenzone der Alpen. Jura und Hügelland*. Stämpflische Verlagshandlung, Friedrich Schulthess Bern.

26. Becker D, Antoine P-O, Engesser B, Hiard F, Hostettler B, Menkveld-Gfeller U, Mennecart B, Scherler L, Berger J-P. 2010 Late Aquitanian mammals from Engehalde (Molasse Basin, Canton Bern, Switzerland). *Annales de Paléontologie* **96**, 95–116. (doi:10.1016/j.annpal.2011.03.001)

27. Menkveld-Gfeller U, Becker D. 2008 Baustelle Zubringer Neufeld: eine neue alte Fossilfundstelle. *Bulletin für angewandte Geologie* **13**, 107–112.

28. Bouvrain G, de Bonis L. 1999 Suoidea du Miocène inférieur de Laugnac (Lot-et-Garonne, France). *Paläontol Z* **73**, 167–178. (doi:10.1007/BF02987990)

29. Morales J *et al.* 1999 Vertebrados continentales del Terciario de la Cuenca de Loranca (Provincia de Cuenca). *La huella del pasado: fósiles de Castilla-La Mancha* , 235–260.

30. Dehm R. 1950 *Die Raubtiere aus dem Mittel-Miocän (Burdigalium) von Wintershof-West bei Eichstätt in Bayern*. München: Verlag der Bayerischen Akademie der Wissenschaften.
